# Synergistic neuroprotective and cognitive-enhancing effects of Walnut Peptide and Theanine in human brain organoid and mouse stress models

**DOI:** 10.1101/2025.02.26.640260

**Authors:** Qixing Zhong, Qinxi Li, Xiuzhen Jia, Linxue Hu, Yingqian Zhang, Jingyang Zu, Yao He, Xiaojie Li, Yu Wang, Haotian Feng, Jingyu Hao, Zifu Zhao, Jian He, Zhihui Zhong

**Author notes:** Corresponding author **Zhihui Zhong** Laboratory of Neurological Disease Modeling and Translational Research, West China Hospital, Sichuan University, No. 37 Guoxue Alley, Wuhou District, Chengdu, 610041, China., **Jian He** Inner Mongolia Dairy Technology Research Institute Co. Ltd.; Yili Innovation Center, Inner Mongolia Yili Industrial Group Co., Ltd., Hohhot, 010110, China. These authors contributed equally to this study (Qixing Zhong, Qinxi Li, Xiuzhen Jia).

## Abstract

Stress is a prevalent mental health concern emerging predominantly in late adolescence or early adulthood. Since 2007, the Food and Drug Administration (FDA) has not approved any novel anxiolytic pharmaceuticals, fueling interest in nutritional supplements as alternative therapies for stress management. Building on prior zebrafish research, this study investigates the synergistic effects of Theanine (*Th*) and Walnut Peptide (*WP*) on stress mitigation and cognitive enhancement. Utilizing the human brain organoid stress (BO-stress) model, *WP*+ *Th* were observed to reduce stress and regulate expression of neurotransmitters, including Gamma-Aminobutyric Acid (GABA), serotonin (5-HT), dopamine (DA), and acetylcholine (Ach), as well as brain-derived neurotrophic factor (BDNF) and serotonin transporter (SERT). Subsequent *in vivo* study using C57BL/6J mouse-stress model demonstrated that the treatment (*Th* 85 mg/mL + *WP* 200 mg/mL), or administrated with vehicles, significantly improved their performance in stress and cognitive assessments, partially normalized neurotransmitter imbalances by modulating SERT and BDNF expression. These findings highlighted the potential of using *WP* + *Th*, particularly when delivered with vehicles (eg: powder/yogurt/milk), as a novel combined therapeutic approach for stress management and cognitive enhancement. We performed a correlation analysis between BO-stress model and mouse-stress model, revealing a alignement in the SERT levels.In addition, SERT was highly correlated with other markers in the mouse hippocampus and may represent a key target for modulating the balance between stress and cognition.

**Significance Statement:** This study established an innovative human brain organoid-stress model and, in conjunction with mouse-stress model, elucidated the synergistic actions of Theanine and Walnut Peptide in mitigating stress and augmenting cognitive function. Their beneficial effects were mediated through the regulation of SERT and BDNF, presenting a promising non-pharmacological avenue for mental health care. In addition, species differences between humans and mice were also examined through correlation analysis of brain organoids and mouse models.

**Graphic abstract:** 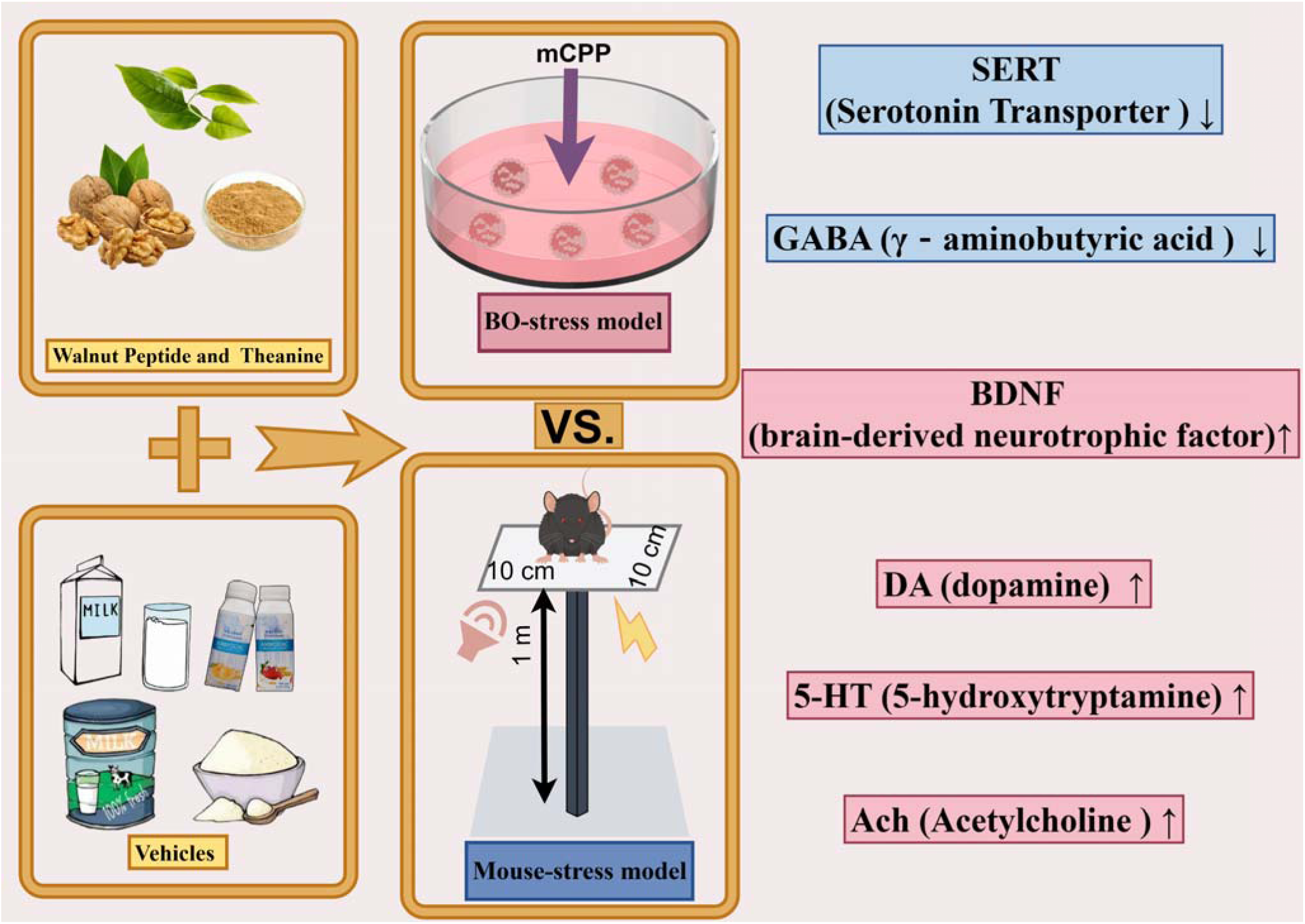

## Introduction

Stress is defined by amplified sense of apprehension or fear in anticipation of potential future threats, whether actual or perceived. Stress-related disorders represent the most common category of mental illness^1^.

Due to their high prevalence, chronicity, and frequent comorbidity with other disorders, the World Health Organization (WHO) has identified stress disorders as the 9^th^ leading cause of disability globally^2^. In addition to emotional manifestations, stress is implicated in neurobiological damage, contributing to conditions such as depression and dementia^3^. A consistent observation in individuals experiencing stress is a deficit in working cognitive. Notably, regardless of the severity of symptoms, those with clinically significant stress scores demonstrate impairments in working cognitive function^4^.

Serotonergic neurons that arise from the brainstem reside within the midbrain’s dorsal periaqueductal gray, with axonal projections extending to various higher-order brain centers, including the prefrontal cortex, hippocampus, and amygdala-regions intimately associated with stress responses^5^. Upon release from the presynaptic neuron, 5-hydroxytryptamine is sequestered by the serotonin transporter (SERT) present in both the presynaptic neuron and neighboring glial cells^6,7^. Beyond serotonergic neurons, stress is capable of modulating the expression of brain-derived neurotrophic factor (BDNF), a key mediator of hippocampal neurogenesis. The hippocampus is integral to learning and cognitive processes. In human context, reduced BDNF expression was reported to link with depression^8,9^. Consequently, the discovery of novel therapeutic agents that concurrently modulate serotonergic neurons and impact BDNF expression represents a priority in our research endeavors.

While pharmacological interventions, such as using antidepressants and benzodiazepines, are available to address stress, their effectiveness varies and may pose a risk of dependency. Since 2007, the US Food and Drug Administration (FDA) has not approved any new anxiolytic agents^10^, underscoring the urgent need for further research to elucidate the mechanisms of stress, as well as to develop safer, more effective therapeutic options^11^. Meanwhile, food and nutrients are gaining recognition as potential alternatives for the management of stress, particularly when delivered with vehicles such as milk and yogurt^12,13^. This natural, safe, and cost-effective strategy offers a manageable option for individuals, potentially mitigating the risks associated with medicines.

Researches have demonstrated that Walnut Peptide (*WP*) enhance cognitive and improve sleep quality in both mice study and clinical trials^14,15^. Similarly, the non-protein amino acid theanine (*Th*), which is prevalent in tea leaves and shares structural similarities with glutamate and glutamine, has also been demonstrated to enhance stress response, sleep quality and cognitive function^16^. Our previous study also revealed that their combination (*WP* + *Th*), exerted significant effects in zebrafish, including antistress, antioxidant, neuroprotective and cognitive enhancement benefits^17^. The combination proved to be more effective than using *WP* or *Th* alone, while also reducing production costs^18^. Furthermore, this combination demonstrated anti-stress and cognitive enhancement effects in stressed mice^19^.

To verify the roles of *WP* and *Th*, we conducted more studies using brain organoids (BOs) derived from human embryonic stem cells (hESCs): We found that *WP+Th* reduced SERT and upregulated BDNF expresisns in BO-stress model. To further validate their anti-stress and cognitive-enhancing effects, we established the mouse-stress model using elevated open platforms (EOP) and physical stressors. Over 30 days, mice were administered with various vehicles containing *W+Th* and their stress levels or cognitive performance were evaluated through a series of behavioral assessments. Furthermore, the levels of neurotransmitters, corticosterone, and neurotrophic factors in the mouse hippocampus (HPC) and prefrontal cortex (PFC) were also measured.

## Materials and methods

### BO-stress modeling and evaluation

HESCs (H9, Beina Biotechnology Co., LTD., China) were cultured with mTeSR1™ medium (Stemcell Technologies, #85850, Canada) and seeded at 9000 cells/well into U-Bottom 96-well ultra-low attachment culture plates (Qingdao AMA CO., LTD, #WP96-6CCUSH, China). The seeding medium was supplemented with 10 µM Y-27632 (Sigma-Aldrich, #Y0503, USA) in embryoid body (EB) formation medium which was refreshed every 2 days. After 5 days, the EBs were transferred to 24-well ultra-low attachment culture plates (Corning Incorporated, #3524, USA) containing induction medium. After 2-day culturing, the EBs were coated with liquid Matrigel^®^ (Corning, #354234, USA). Subsequently, they were transferred to pre-treated 6-well plates (NEST, #703001, USA) with 12-16 organoids per well. The embedded BOs were maintained in expansion medium for 3 days. On day 10, the BOs were transferred into mature medium and placed on orbital shaker (JOANLAB, #OS-20, China) at 70 rpm. The culture medium was then refreshed every 4 days (workflow is shown in **Figure 1A**).

**Figure 1.**
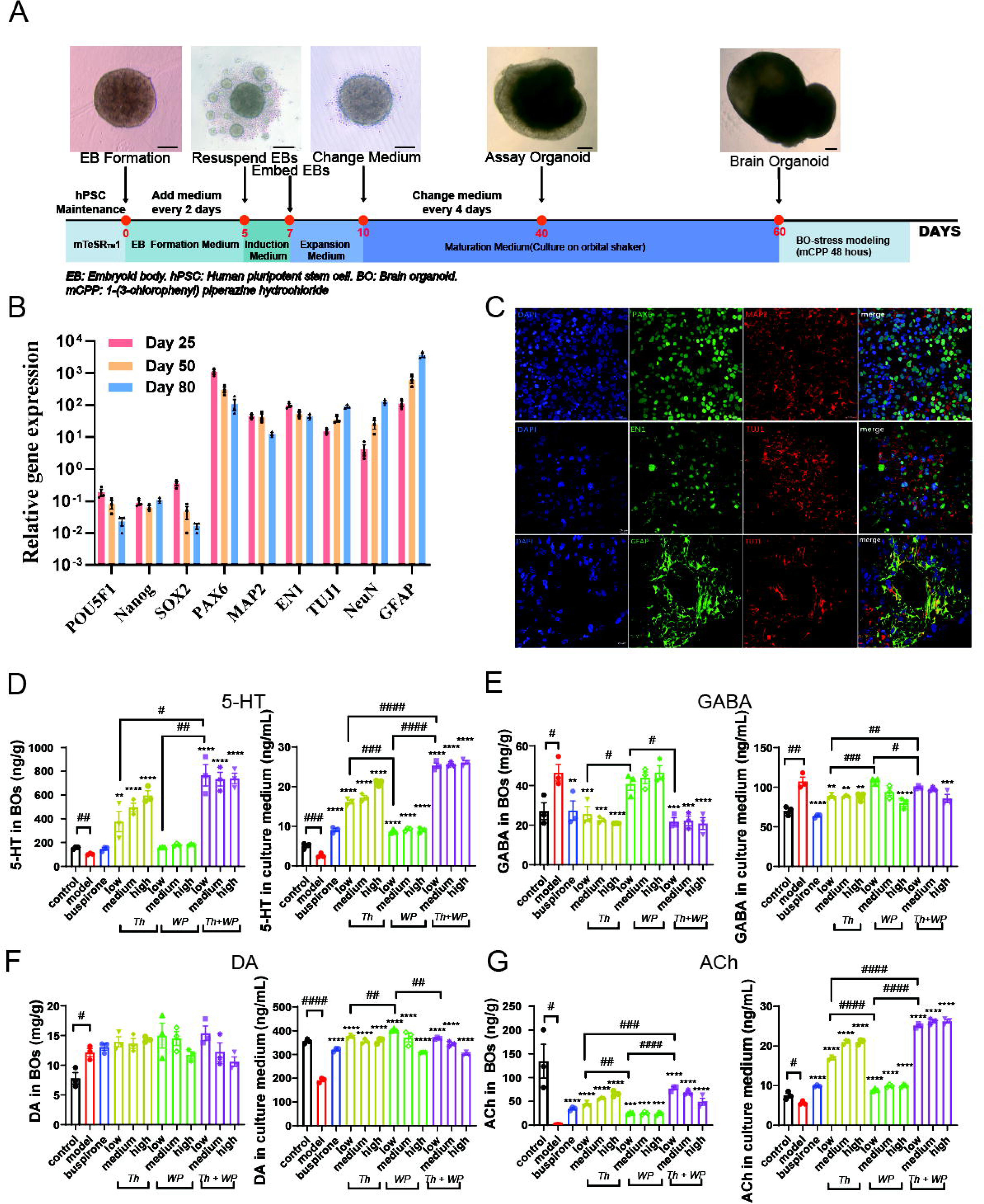
*Th+WP* regulated neurotransmitter in BO-stress model. (**A**) The workflow of BO establishment and stress modelling. Scale bar = 100 μm. (**B**) RT-qPCR analysis of gene expression relative to GAPDH and H9 ESCs. Data were presented as mean ± SEM, n = 3. (**C**) IF analysis of BOs on day 60, DAPI (blue), PAX6, EN1,GFAP (green) and MAP2, TUJ1(red). The gene expression level was normalized to the expression of GAPDH and H9. Scale bar = 20 μm. The concentrations of 5-HT (**D**), GABA (**E**), DA (**F**), and ACh (**G**) in the BOs and medium were measured by LC-MS/MS. The data were presented as mean ± SEM, n = 3, in each group (\**P* < 0.05, \*\**P* < 0.01, \*\*\**P* < 0.001, \*\*\*\**P* < 0.01 vs. model group. #*P* < 0.05, ##*P* < 0.01, ###*P* < 0.01, ####*P* < 0.0001).

1-(3-chlorophenyl) piperazine hydrochloride (mCPP, #CDS002717, Sigma, USA) was used to induce the BO-stress model. To explore the optimal condition of mCPP, we leveraged stress-induction concentration from previous zebrafish study^17^, and systematically varied the mCPP concentrations across a range of 20 to 10,000 µg/mL. The duration of exposure was tested from 1 to 48 hours. The efficacy of the stress model was determined by measuring ATP levels (Promega, #G9681, USA) (**Supplementary** Figure 1A,B) and cortisol using ELISA kit (Vankovic Biotechnologies, #F0770-A, China) (**Supplementary** Figure 1C,D) and serotonin (5-HT, Vankovic Biotechnologies, #F111342-A, China) (**Supplementary** Figure 1E,F), in both the BOs and the culture medium. After that, the compound was thoroughly washed away.

Subsequently, BOs (day 60) were randomly allocated into 6-well plates and classified into 12 groups (n = 3): control, model, buspirone, *Th*-low, *Th*-medium, *Th*-high, *WP*-low, *WP*-medium, *WP*-high, (*Th+WP*)- low, (*Th+WP*)-medium, and (*Th+WP*)-high. BOs of the control were maintained in culture medium. The remaining 11 groups were subjected to 48-hour exposure to 1 mL of culture medium containing mCPP (2.4 mg/mL) for modeling. The buspirone (#33386-08-2, Sigma, USA) was added at 20 mg/kg in 1 mL of culture medium. Dosages of the remaining groups are reported in **Table 1** below. After 24-hour treatment, the culture medium and BOs from each group were collected for further analysis.

**Table 1.** The sample information of BO experiments.

| Group | Name | Origin | Human dose (mg/day) | BO dose ($\mu\text{g/mL/day}$) |
| --- | --- | --- | --- | --- |
| **Control** | N.A | N.A | N.A | N.A |
| **Model** | mCPP | Sigma Chemical Co | N.A | N.A |
| **Buspirone** | Buspirone Hydrochloride | Sigma Chemical Co | 20 | 3.33 |
| ***Th*-low** | The natural amino acid in green tea | Shanghai Novanat Co., Ltd | 200 | 33.3 |
| ***Th*-medium** |  |  | 300 | 49.9 |
| ***Th*-high** |  |  | 400 | 66.6 |
| ***WP*-low** | The plant extract, bioactive peptide extracted from the protein of walnut residues | Sinphar group Co., Ltd | 85 | 14.2 |
| ***WP*-medium** |  |  | 170 | 28.4 |
| ***WP*-high** |  |  | 400 | 66.6 |
| **Combinations** | (*Th*+*WP*)-low | N.A | 200 + 85 | 33.3 + 14.2 |
|  | (*Th*+*WP*)-medium | N.A | 200 + 170 | 33.3 + 28.4 |
|  | *(Th+WP)-high* | N.A | 400 + 170 | 66.6 + 28.4 |

Notes: N.A: not applicable. Data obtained from FDA draft guidelines<sup>20</sup>. The works of Zhao et al<sup>17</sup> and Rita Fior et al<sup>21</sup> provide insights into the conversion methodology between human dosage and zebrafish dosage, given that the volume of BOs closely resembles that of zebrafish..

### LC-MS/MS analysis

LC-MS/MS (Waters HPLC, AB API 5000 LC/MS/MS) and Analyst^®^ (Gerstel GmbH & Co. KG) software was used to detect 5-hydroxytryptamine (5-HT), Acetylcholine (ACh), γ-aminobutyric acid (GABA), and dopamine (DA) in the hippocampus (HPC) and prefrontal cortex (PFC) of mice brain. A column (50 mm ×2.1 mm i.d.; particle size, 1.7 μm; Waters, Milford, USA) was used at 40 [. The chromatographic gradient is shown in **Supplementary Table 1**. The flow rate was 0.60 mL/min, with injection volume of 5 μL, dexamethasone and verapamil were used as internal standards.

### Immnuoflourences (IF) analysis

BO sections underwent dewaxing to the aqueous phase, succeeded by antigen retrieval employing a thermal epitope retrieval technique. These protocols were aligned with the procedures for fixing and staining the PFC of mice. Following perfusion with normal saline and 4% paraformaldehyde (PFA) solution, brain tissues were harvested and immersed in 4% PFA for 24-hour fixation. Dehydration ensued in a 30% sucrose solution in 0.01 M PBS for 48 hours. Serial brain sections of 25 µm thickness were procured using a cryostat microtome (Leica CM1860UV, Germany). Subsequent to 30-minute permeabilization with 0.3% Triton and 1-hour blocking with 5% BSA, the sections were incubated with primary antibodies (as detailed in **Supplementary Table 2**). DAPI (1:1000, Thermo Fisher Scientific, USA) was then applied for 10- minute incubation at room temperature. Post-staining, sections were rinsed thrice with PBS (5-minute each). Sections were mounted with an anti-fade reagent (Solarbio, China) and imaged using an inverted confocal microscope (Andor Dragonfly 200, UK). To counteract fluorescence decay, sections were treated with an anti-quenching reagent (Solarbio, China) prior to confocal microscopy analysis. Image analysis was facilitated by Imaris Viewer software (Oxford Instruments, UK), while quantification of fluorescence-expressing regions within IF images was conducted using ImageJ software. The relative protein expression levels across images were determined by the following formula:

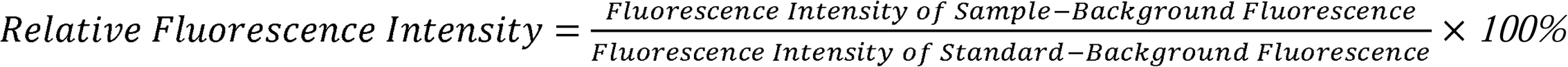

### RNA extraction and RT-qPCR analysis

TRIzol reagent kit (#15596026, Invitrogen) was used to isolate the total RNA. RNA purity and yield were measured using NanoDrop 2000 spectrophotometer (Thermo Scientific, USA). RNA integrity was evaluated using Agilent 2100 Bioanalyzer (Agilent Technologies, USA). Adhering to the protocol provided by the manufacturer, the VAHTS Universal V6 RNA-seq Library Prep kit (#NR604-01, Vazyme, China) was utilized to prepare the transcriptome libraries. Thereafter, the sequencing libraries underwent sequencing using the NovaSeq 6000 System (Illumina). RNA was reverse transcribed into cDNA using HiScript III RT SuperMix for qPCR (+gDNA wiper) (Vazyme Bioinformatics Technology Co., Ltd, #R323, China) and ChamQ Universal SYBR qPCR Master Mix (Vazyme Bioinformatics Technology Co., Ltd, #Q711, China). The primer sequences used are listed in **Supplementary Table 3**. The Bio-Rad CFX Manager 3.1 (Bio-Rad Technology Co., Ltd, CFX Connect, USA) was employed for RT-PCR analysis with the manufacturer-recommended parameters.

### Animals

50 male and 50 female C57BL/6J strain mice, aged 6-8 weeks, were procured from Charles River Laboratories Animal Technology Co., Ltd (Beijing, China). These mice were housed under specific pathogen-free (SPF) conditions, following 12-hour light-dark cycles, and provided standard chow and water sourced from Beijing Keao Xieli Feed Co. The mice (4 mice per cage, animal production license number was SYXK (Chuan) 2019-215) were maintained at the Sichuan Junhui Biotech facilities. All experimental procedures adhered to the guidelines set forth by the Association for Assessment and Accreditation of Laboratory Animal Care (AAALAC) and were approved by the Institutional Animal Care and Use Committee (IACUC) of the West China Hospital, Sichuan University (Approval No. 2019194A).

### Mouse-stress modelling and grouping

Based on previous study^22^, we established mouse-stress model using EOP and physical stress. Mice were exposed to clear square Plexiglas board (10 cm × 10 cm) at 1 metre for 1 hour/day for 14 days (**Supplementary** Figure 2A). Control mice were kept under standard conditions without EOP exposure, while maintaining uniform living conditions. On day 1, mice were randomized by weight and divided into 10 groups (10 mice in each group and their body weight is shown in **Supplementary** Figure 2B).

Body weight, food intake, and coat state score were evaluated every 7 days during 10-12 a.m. The coat state was assessed in seven areas of the mice, including the head, neck, dorsal area, ventral area, tail, anterior claw, and hind claw. A coat with regular, soft hair was recorded as 0 points, while an unwell- groomed coat with dark hair color was assessed as 1 point for each area^23^. Buspirone hydrochloride was dissolved in saline and administered intraperitoneally for 30 days (2 mg/kg) daily^24^. This product comprises 90% oligopeptide, with the majority falling within the range of < 1000 Da. Details of the dosing protocols for mice are listed in **Table 2** below.

**Table 2.**
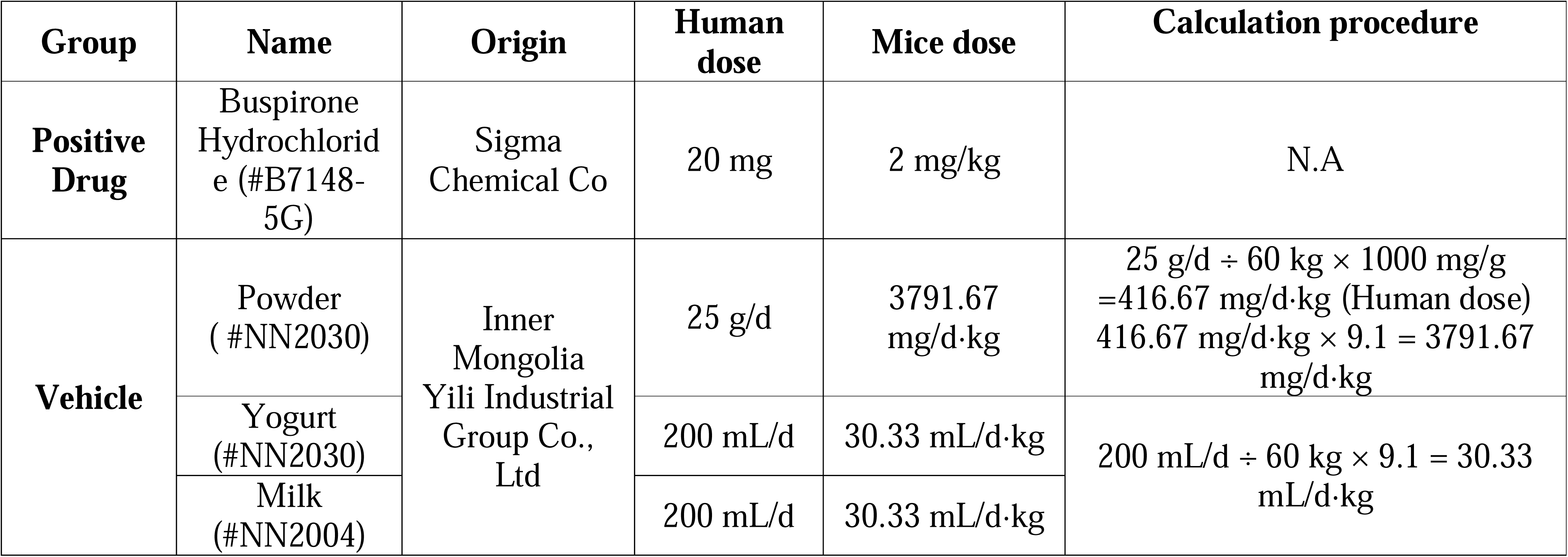
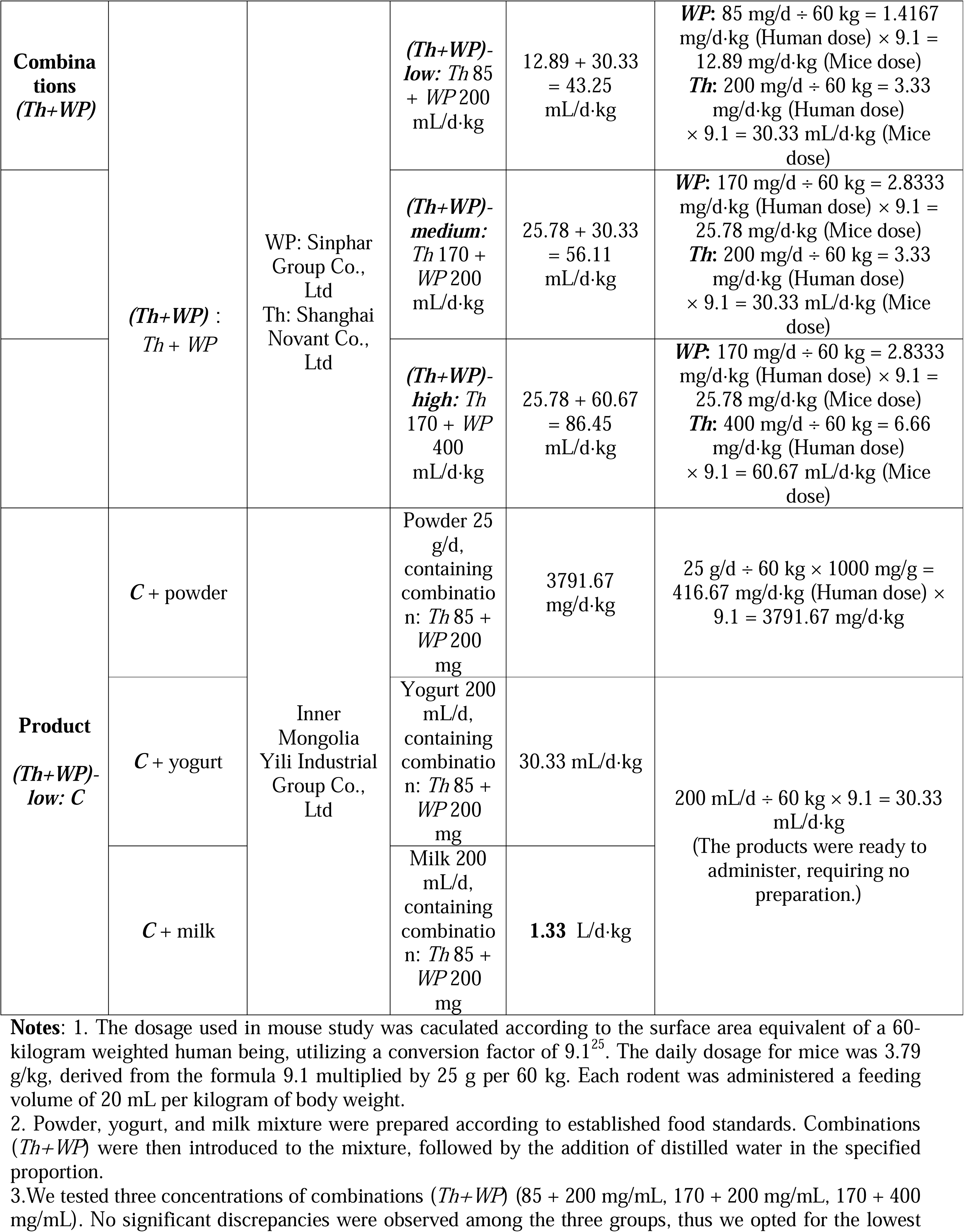
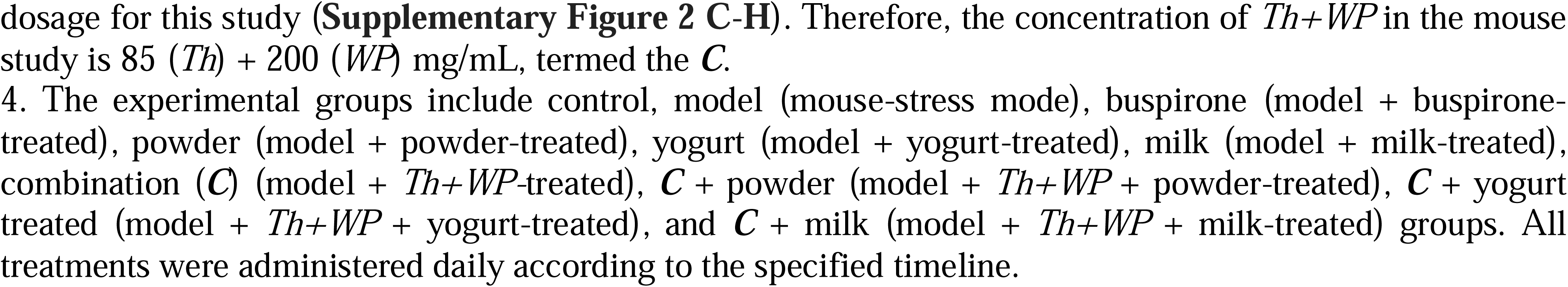
The sample names, origins and concentrations in mouse experiments.

### Open-field test (OFT)

As described previously, each mouse was gently introduced into a 50 × 50 × 50 cemtimeters (cm) open field arena,^26^ where it was allowed to explore freely for 5 minutes under dim illumination (50 lx). The total distance traveled (in cm) and the frequency of entries into the central zone (25 × 25 cm) were recorded using the Topscan Package (Clever Sys Inc., USA).

### Elevated plus-maze (EPM) test

The EPM consisted of two open arms and two closed arms. Each arm was 30 cm in length and 5 cm in width, with 20 cm high walls in the closed arms. The EPM was elevated 60 cm above the ground, with a 90° angle between each arm. Mice were initially placed in the central area facing an open arm and allowed to explore for 5 minutes^27^.

### Light-dark box (LDB) test

As previously described^28^, stress responses were evaluated using light/dark box paradigm. Mice were housed in a two-chamber Plexiglas apparatus (18 × 12 × 12 cm) featuring a central opening (5 × 5 cm). The mice were positioned in the light chamber facing the dark chamber and their exploratory behaviors were recorded for 10 minutes following their passage through the opening door.

### Novel object recognition (NOR) test

Recognition was tested using NOR test as previously described^29^. During learning trial, mice were exposed to two indistinguishable objects within an open field box for a duration of 10 minutes. Following one-hour interval, the test trial was commenced. In the course of the test trial, one new object and one identical object were introduced to replace the previous objects, and the mice engaged in exploration for a period of 5 minutes.

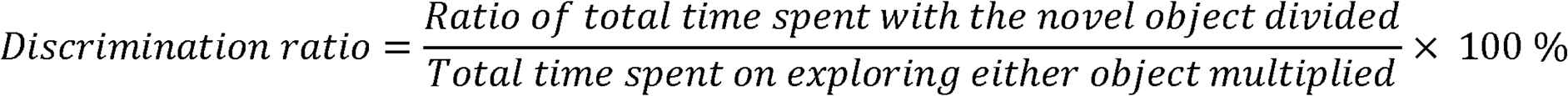

### Passive avoidance test (PAT)

The PAT was used in a light/dark shuttle box from day 36-40, followed reported method with modifications^30^. On day 1, mice engaged in 2-minutes exploration of both the dark and bright compartments. Subsequently, on day 2, mice were placed within the illuminated box again, while subjected to an electric shock of 0.6 mA for 2 seconds. On the test day, mice were placed exclusively in the bright compartment and allowed to explore for 2 minutes. During this time, no electric shock was administered when it entered the dark compartment.

### Enzyme-linked immunosorbent assay (ELISA)

Mice were sacrificed by exposure to carbon dioxide during 7:00-8:00 a.m., coinciding with the blood collection period. The levels of corticosterone and ACTH in serum were ascertained using ELISA kits (TECAN, #RE52211, Germany) and (abcam, #ab263880, United Kingdom), respectively.

### Sample preparation of mouse brain tissue

After the complete dissection of the mice, their brain were bilaterally segregated, dividing into left and right hemispheres. The entire left hemisphere was exclusively utilized for the detection of Iba1 immunohistochemistry. Subsequently, the right hemisphere was promptly placed on ice to facilitate the extraction of the right HPC and the PFC. The PFC region of the right hemisphere was subjected to LC-MS/MS analysis. The HPC of the same hemisphere was prepared for RT-qPCR analysis.

### Statistical analysis

Statistical analysis was performed using SPSS 26.0 software (IBM Ltd, UK). Data with normal distributions were presented as mean ± SEM. Differences between control and model were assessed using unpaired t-test. Differences among the model, buspirone, and each treatment group were compared using one-way ANOVA followed by post hoc Tukey Dunnett’s multiple comparison test. Repeated measures ANOVA was used to evaluate differences in body weight, food intake, and coat state score between groups. The significance of correlations between each effect was determined using Pearson’s correlation. A significance level of *P* < 0.05 was used for all analyses.

## Result

### Th+WP regulated neurotransmitter in BO-stress model

We firsly utilized BOs derived from hESCs to establish BO-stress model (work flow is shown in **Figure 1A**). During BO establishment, we analyzed the gene expression throughout the differentiation. Results in **Figure 1B** showed a low expression pluripotency marker POU5F1, Nanog or SOX2. A pile up-expression of neuronal markers TUJ1, NeuN as well as marker of astrocytes (GFAP). We futher conducted IF analysis to confirm the formation of neurons and astrocytes in BOs (**Figure 1C**). Subsequently, we utilized BOs to investigate stress by introducing mCPP at various concentrations. In ATP assays, BOs showed no significant changes in response to variations in the timing and concentration of mCPP addition. (**Supplementary** Figure 1A**, B**). In addition, 48-hour addition of mCPP (at 2.4 mg/mL) resulted higher cortisol concentration than the control (**Supplementary** Figure 1C**, D**). For 5-HT, we noted a gradual decline in concentration upon mCPP addition, reaching a stable level after 48 hours. Futher concentration test at this time point showed a lower 5-HT level from 2.4 mg/mL (**Supplementary** Figure 1E**, F**). Based on above results, we selected 48 hours treatment of mCPP at 2.4 mg/mL for BO stress modelling. Subsequently, the BO-stress models were treated with buspiroine, *Th*, *WP* and *Th+WP* (with low, medium, high concentrations, reported in the method section and listed in **Table 1**), and the concentrations of neurotransmitters in both the BOs and their culture medium were measured. LC-MS/MS analysis demonstrated a significant 5-HT level elevation in BOs and their culture medium following treatment with *Th* at low, medium, and high doses, as well as in combination with *WP* at similar dose. Notably, 5-HT levels in the *WP* group were significantly lower that in the *Th*+*WP* treatment groups (**Figure 1D**, both in BOs *P* < 0.05 and the culture medium *P* < 0.0001). Compared to the model, GABA expression in BOs were markedly reduecd by *Th* treatment at low, medium, and high doses, as well as with combined *Th+WP* at equivalent dosages. Intriguingly, GABA levels in *WP* group were considerably elevated compared to those in *Th+WP* group in BOs (*P* < 0.05). In the culture medium, the *WP*-low group displayed significantly elevated levels compared with the *Th*-low group (*P* < 0.001) and the *Th+WP*-low group (*P* < 0.05) (**Figure 1E**). In BOs, DA levels were significantly lower in the control group than in the model (*P* < 0.05), with no significant differences detected between the model group and other groups. In the culture medium, the model group displayed significantly lower levels compared with other groups. Notably, the *WP*-low group exhibited significantly higher levels than both the *Th*-low and *Th+WP*-low groups (*P* < 0.01, **Figure 1F**). Regarding Ach, the model group showed significantly lower levels compared to the *Th* (low, medium, and high doses) and *Th*+*WP* (low, medium, and high doses) groups, with significant differences observed between the *WP* and *Th*+*WP* groups. In the culture medium, the model group exhibited a significantly lower concentration of Ach compared with all other groups. Notably, the low-dose treatment in the *Th* group resulted in a significantly higher concentration of Ach compared with the *WP* group (*P* < 0.01), and the *Th+WP* group exhibited a significantly higher Ach concentration than the *Th* group alone (*P* < 0.001). This suggests that the *Th* group is the primary driver of Ach elevation, with the *Th+WP* combination yielding a more pronounced effect on Ach concentration (**Figure 1G**). Collectively, the *Th+WP* administration resulted in elevated 5-HT and Ach levels in BOs, predominantly attributed to the action of *Th*. Furthermore, this addition led to a reduction in GABA concentrations in BOs, largely influenced by the presence of *WP*. Since there were no significant differences in 5-HT, GABA, and ACh levels across the low, medium, and high dose groups for *Th*, *WP* or *Th+WP*, we selected the lowest concentration of *Th* (85 mg/mL), *WP* (200 mg/mL), and *Th+WP* (85 + 200 mg/mL) for further tests.

### Th + WP regulated SERT and BDNF in BO-stress model

SERT is a key membrane protein that facilitates the re-uptake of serotonin from the synaptic cleft into the presynaptic neuron. This process is essential for regulating serotonin levels and is a primary target for selective serotonin re-uptake inhibitors (SSRIs), which was used to treat mood disorders^31^. In our study we found that, in the BO-stress model, SERT expression was elevated compared to control (*P* < 0.0001), and was then reduced by the addition of buspirone (*P* < 0.05). Similarly, the treatment with *Th, WP*, or *Th+WP* lowered SERT expression levels, with *Th+WP* being the most effective (**Figure 2A,B**). Compared to the model, similar reduction of the SERT mRNA expression in the buspirone, *Th*, *WP* and *Th +WP* group were observed. The mRNA expression of SERT in the *WP* group was significantly higher than that in the *Th* group and the *Th+WP* group (*P* < 0.001, respectively) (**Figure 2B**).

**Figure 2.**
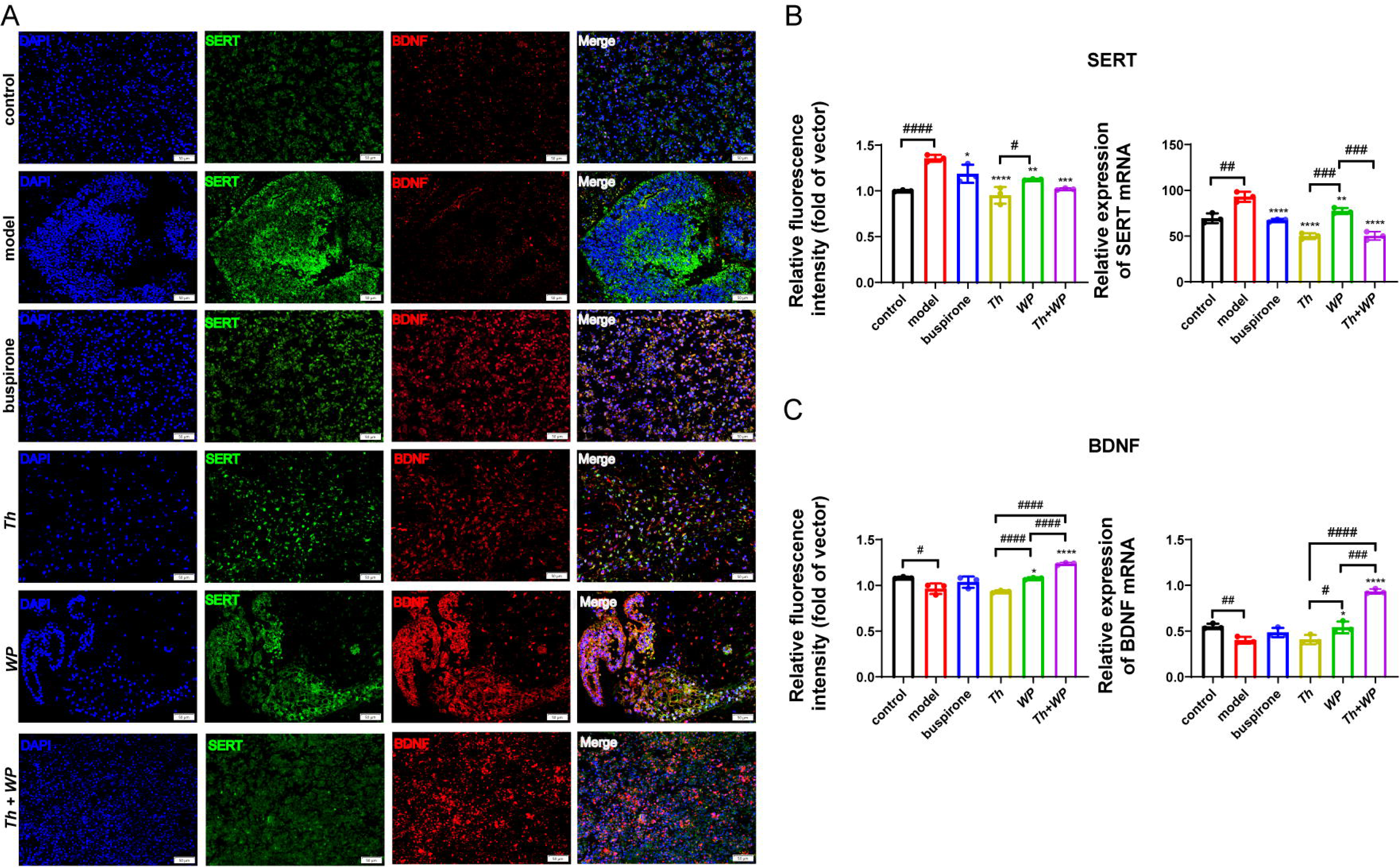
*Th + WP* regulated SERT and BDNF in BO-stress model. (**A**) IF analysis of BOs on day 60, DAPI (blue), SERT (green) and BDNF (red). Scale bar = 20 μm. (**B**) Left: Quantification of SERT-associated relative fluorescence intensity. Data were represented as mean ± SEM, n = 3. Right: RT-qPCR analysis of SERT normalized to GAPDH and H9 ESCs. Data were presented as mean ± SEM, n = 3. (**C**) Left: Quantification of BDNF-associated relative fluorescence intensity were presented as mean ± SEM, n = 3. Right: RT-qPCR analysis of BDNF normalized to GAPDH and H9 ESCs. Data were presented as mean ± SEM, n = 3. Significance was defined as *P* < 0.05 (\**P* < 0.05, \*\**P* < 0.01, \*\*\**P* < 0.001, \*\*\*\**P* < 0.01 vs. model group. #*P* < 0.05, ##*P* < 0.01, ###*P* < 0.01).

BDNF is crucial in depression’s pathophysiology^32^. Antidepressants like SSRIs boost BDNF, promoting synaptic plasticity, neurogenesis, and neuronal survival via TrkB receptors^33^. In our study, compare to the control, the model exhibited lower expression of BDNF (*P* < 0.05). The relative fluorescence intensity and mRNA expression of BDNF was up-regulated by the administration of *Th, WP and Th+WP,* with *Th*+*WP* being the most effective. At the mRNA level, the *Th+WP* group exhibited significantly higher BDNF expression than either the *Th* group (*P* < 0.0001) or the *WP* (*P* < 0.001) group, with the *WP* group also showing significantly elevated BDNF expression compared with the *Th* group (*P* < 0.05) (**Figure 2C**).

The results indicate that *Th+WP* can both reduce the expression of SERT and increase the expression of BDNF.

### The C showed anti-stress effect in mouse-stress model

In order to substantiate the antidepressant efficacy of *Th+WP*, we employed the elevated open platforms (EOP) modeling technique to develop a mouse-stress model (**Supplementary** Figure 2A). In pursuit of experimental rigor, we assessed the effects of low, medium, and high doses of *Th+WP* using the mouse-stress model, revealing no significant differences (**Supplementary** Figure 2C-H). Therefore, in the following experiments, we continue to investigate whether the *Th*+*WP* (85 + 200 mg/mL, termed as the ***C***) alleviate stress and improve cognitive abilities in mouse-stress model. Furthermore, we expanded our study to investigate the effect of administrating ***C*** with vehciles: powdered milk (referred to as powder), liquid milk (referred to as milk), and yogurt^34,35^. During modelling, stress and cognitive levels in the mouse- stress model were evaluated via behavioral experiments conducted at specific time points (workflow is shown in **Figure 3A**). Assessments of body weight, food consumption, and coat condition were conducted to evaluate the physiological status across groups. While the model group exhibited elevated coat condition scores, these differences were not statistically significant, as illustrated in **Supplementary** Figure 3A-C. Outcomes of a 14-day stress-induced mouse model assessed via OFT was preseted in **Supplementary** Figure 3D-F. These results confirm the successful establishment of EOP-induced stress model in mice and indicate that treatment with ***C***, or supplemented with milk or yogurt, significantly mitigated stress responses. In the OFT, only the model exhibited a significant decrease in center entries compared to controls (*P* = 0.001, **Figure 3B-D**). Metrics such as center entries and travel distance primarily reflect horizontal movement, they may not adequately capture vertical activity. Therefore, rearing behaviors thus serve as a more sensitive measure of stress in rodent behavioral studies^36^. The minimum rearing time they appeared in the model group were significantly less than the control (*P* < 0.01) or the buspirone (*P* < 0.05). The ***C*** increased rearing times compared to model (*P* < 0.0001), ***C*** + powder (*P* < 0.01), and ***C*** + yogurt (*P* < 0.01) (**Figure 3C, D**).

**Figure 3.**
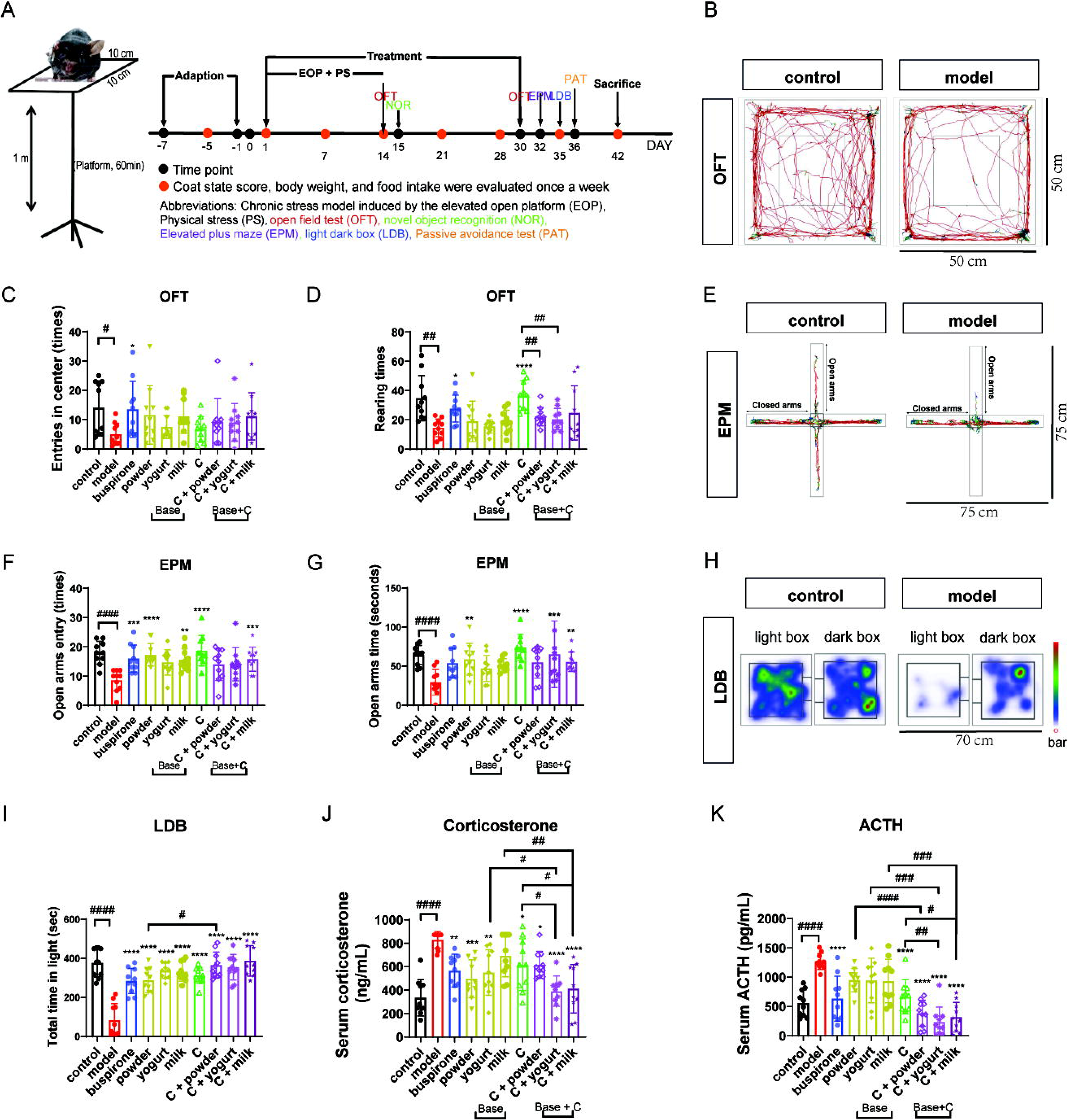
The *C* showed anti-stress effect in mouse-stress model. Abbreviations: Chronic stress model induced by the elevated open platform (EOP), Physical stress (PS), open field test (OFT), novel object recognition (NOR), elevated plus maze (EPM), light dark box (LDB), passive avoidance test (PAT). (**A**) Left: EOP photos: Transparent 10 cm square platform at 1m height, mice exposed for 60 min. Right: EOP timeline: 7-day acclimation, EOP and treatments from day 1-30, OFT and NOR on day 14, OFT, EPM, LDB, AT on days 30, 32, 35, 36. Mice sacrificed on day 42 for tissue and serum collection. **B**-**D:** OFT performed on the day 30. The representative pictures of control and model (**B**), changes of rearing times (**C**), and entries in the center (**D**) by OFT in each group. **E-G**: EPM performed on day 32. Representative pictures of EPM (**E**), changes of open arms entry (**F**), and open arm time (**G**) by EPM in each group. LDB by trajectories on day 35. Representative pictures of LDB (**H**) and changes of total time in light of each group (**I**). **J-K:** Serum concentrations of corticosterone (**J**) and ACTH (**K**) in each group. Data represent mean ± SEM; n = 10 in each group. Significance was defined as *P* < 0.05 (\**P* < 0.05, \*\**P* < 0.01, \*\*\**P* < 0.001, \*\*\*\**P* < 0.01 vs. model group. #*P* < 0.05, ##*P* < 0.01, ###*P* < 0.001, ####*P* < 0.0001).

In the EPM test, time and duration of open arms entry were the least in the model group. The buspirone, milk, ***C***, and ***C*** + powder/yogurt/milk explored open arms much more than the model (**Figure 3E-G**), indicating that ***C*** or supplemented with vehicles could improve the stress state in mice. The LDB test served as the third behavioral assessment for evaluating stress. The model spent less time in the light chamber compared to control (*P* < 0.0001), which was reversed by the treatment with buspirone (*P* < 0.0001). Furthermore, in the light-dark box (LDB) test, the duration for which the ***C***+powder group remained in the light compartment was significantly greater than that of the powder group (*P* < 0.05, **Figure 3H,I**).

Next, we examined changes in stress-associated hormones in the serum. Following EOP stress exposure, a significant elevation in serum corticosterone levels was observed in mice, with a marked distinction between the control and model groups (*P* < 0.0001). The administration of ***C*** + yogurt or ***C*** + milk resulted in a pronounced reduction in corticosterone levels. Furthermore, mice that received yogurt or milk alone exhibited significantly higher corticosterone levels compared to those that received ***C*** + yogurt or ***C*** + milk (**Figure 3J**). Changes in serum ACTH level were observed to be more sensitive than corticosterone: Compare to model, serum ACTH levels in ***C***, and ***C*** + powder/yogurt/milk groups were lower. The ACTH values of group ***C*** were significantly higher than those of ***C*** + yogurt/milk (*P* < 0.01, *P* < 0.05, respectively). Compare to ***C***, administration with vehicles further reduced ACTH expression (**Figure 3K**). In summary, the ***C*** exerts a pronounced effect on alleviating stress-related behavioral indices in mice. Serum corticosterone and ACTH levels further demonstrate that the efficacy of ***C*** is significantly enhanced when administered in combination with milk powder, yogurt, or milk.

### The C enhanced cognitive in the mouse-stress model

Subsequently, we employed NOR and PAT to evaluate cognitive in mouse-stress model. Compared to control, NOR tests conducted on day 15 revealed a significant decrease in the recognition index in the model , indicating that chronic stress induced cognitive impairment. In contrast, treatment with buspirone, ***C***, or ***C*** combined with vehicles enhanced cognitive. Notably, the recognition index of the ***C***+milk group was significantly higher than that of the milk group alone (**Figure 4A,B**). In the PAT, the model exhibited shortest latency to enter the dark box. In contrast, treatment with buspirone, yogurt, ***C***, and ***C*** + vehicles significantly extended the latency to the dark box, highlighting their potential to mitigate stress-induced cognitive impairments. Furthermore, the latency to enter the dark box was markedly increased in the ***C***+powder/yogurt group compare to the powder/yogurt group (**Figure 4C,D**), indicating that ***C***+base is more conducive to cognitive enhancement than the vehicles.

**Figure 4.**
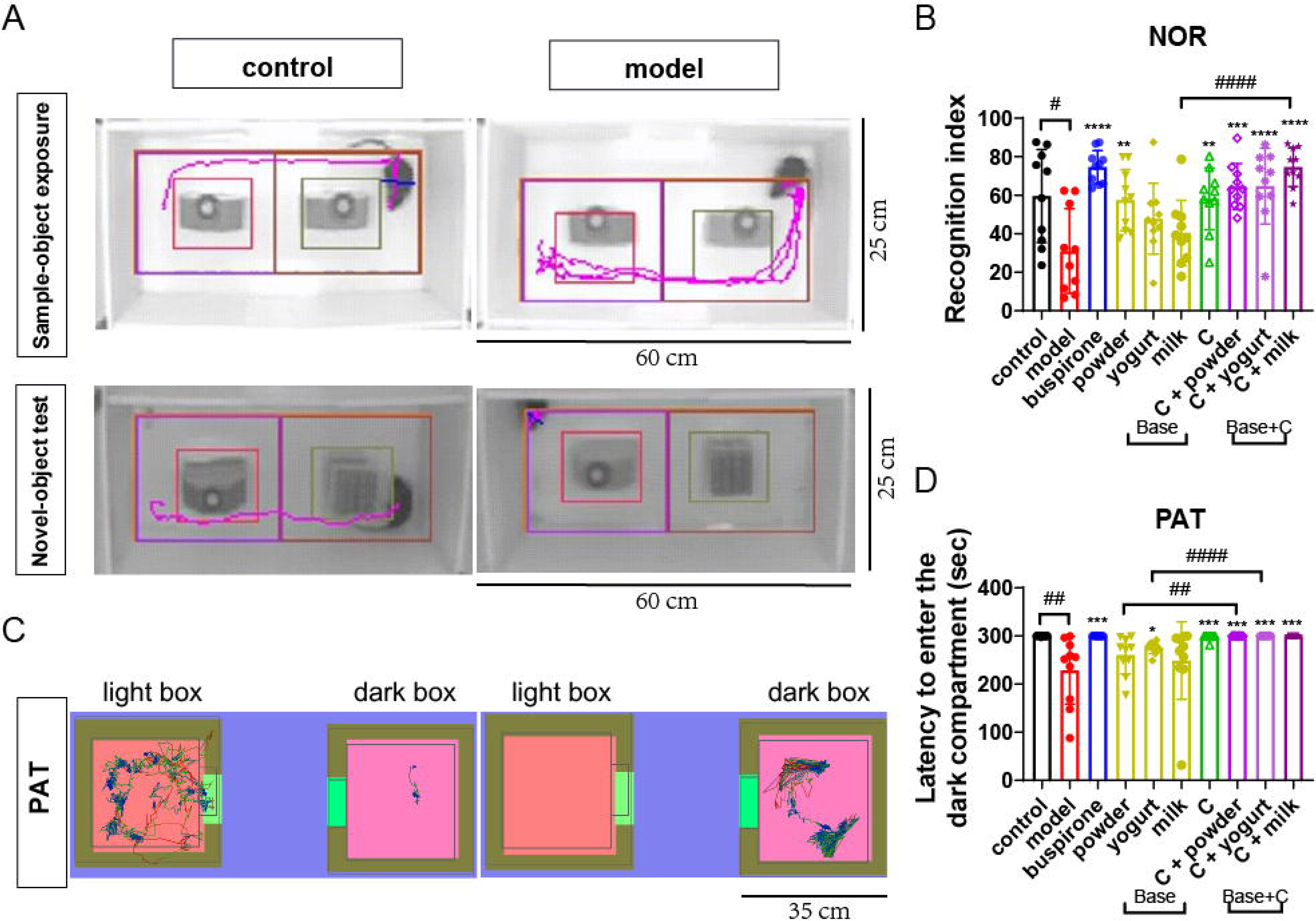
The *C* enhanced cognitive in the mouse-stress model. **A-B:** The NOR test performed on day 15. Representative track images of control and model groups were shown in (**A**). The time spent exploring one of the objects during the familiarization session (left) and the novel object during the test session (right) for the two experimental procedures tested. The recognition index analyzed by NOR in each group (**B**). **C-D:** The passive avoidance test performed onday 36. Representative track images of control and model groups were shown in (**C**). Latency to enter the dark compartment by PAT in each group on day 36 (**D**). Data represent mean ± SEM; n = 10 in each group. Significance was defined as *P* < 0.05 (\**P* < 0.05, \*\**P* < 0.01, \*\*\**P* < 0.001, \*\*\*\**P* < 0.01 vs. model group. #*P* < 0.05, ##*P* < 0.01, ###*P* < 0.01, ####*P* < 0.0001).

### C re-balanced SERT and BDNF expressions in mouse-stress model

SERT primarily regulates the reuptake of 5-HT in the hippocampus (HPC), and its role in modulating emotional and cognitive functions has been widely recognized^37^. The expression and function of SERT in the prefrontal cortex (PFC) are closely associated with the pathogenesis of neuropsychiatric disorders like depression^38^. Research has shown that early-life stress (ELS) can downregulate BDNF expression in the HPC by upregulating the levels of H3K9me2, affecting cognitive functions^39^. Blocking the TrkB receptor in the PFC significantly impairs reversal learning in rats, although its impact on spatial learning ability is relatively limited^40^. Therefore, it was of our interest to investigate their relationships in our mouse-stress models. Firstly, we observed that the expression of SERT increased in mice after stress, showing significant differences compared to the groups treated with ***C***, and ***C*** combined with vehicles. Moreover, the expression of SERT in the ***C*** + powder and ***C*** + milk groups was significantly lower than that in the groups treated with powder or milk alone (**Figure 5A**, left panel). In addition, the PFC, renowned for its susceptibility to internal stressors, is crucial in modulating emotional responses and serves as the principal locus for BDNF’s activity^41^. We showed that, the model group mice exhibited significantly reduced BDNF mRNA expression in the HPC region compared to the control (*P* < 0.05). Following the treatment with ***C*** + vehicles, BDNF expression was elevated and significantly surpassed that of the groups receiving vehicles alone (**Figure 5A**, right panel). In contrast to control, the model exhibited lower 5-HT level (*P* < 0.05). In ***C***, and ***C*** + vehicles, higher 5-HT levels than the model were observed. ***C*** + vehicles had a higher 5-HT concentration than vehicle only (**Figure 5B**, left panel). In contrast, the model group had higher GABA concentrations than control. The GABA levels in the ***C*** + yogurt group were significantly lower than those in the individual yogurt groups (**Figure 5B**, right panel). Similarly, DA concentration was much higher in the model than control. ***C*** *+* yogurt and ***C*** + milk reduced DA concentration compared with model (**Figure 5C**). In the model, the level of Ach was markedly reduced compared to control, ***C***, and ***C*** + vehicles. Notably, the ***C*** + powder/yogurt group exhibited a significant increase in Ach levels relative to the ***C*** group. In addition to the aforementioned analyses, we assessed the PFC and observed outcomes akin to those in the HPC region (**Figure 5D**-**F**). Of particular interest, the ***C*** + powder group exhibited markedly elevated BDNF expression relative to ***C*** and ***C*** + yogurt/milk groups. The group administered powder alone demonstrated significantly reduced BDNF expression when compared to the ***C*** + powder group, suggesting that ***C*** + powder exerts a more pronounced influence on BDNF expression within the PFC region. Previous studies have elucidated the pivotal role of microglia in modulating stress responses within murine models^42^. Therefore, we further explored the effect of ***C*** on microglia. Results showed no significant differences in the number of Iba1- positive microglial cells between control and model, indicating that the elevation in Iba1 expression and coverage was not due to differences in cell number. However, compared to the model, a significant reduction in the number of Iba1positive cells was observed in the ***C*** + powder and ***C*** + yogurt groups (**Supplementary** Figure 4), suggesting a potential modulatory effect of ***C*** + powder and ***C*** + yogurt interventions on microglial reduction. Further investigation, including subsequent microglial phenotyping, is essential to establish a definitive conclusion.

**Figure 5.**
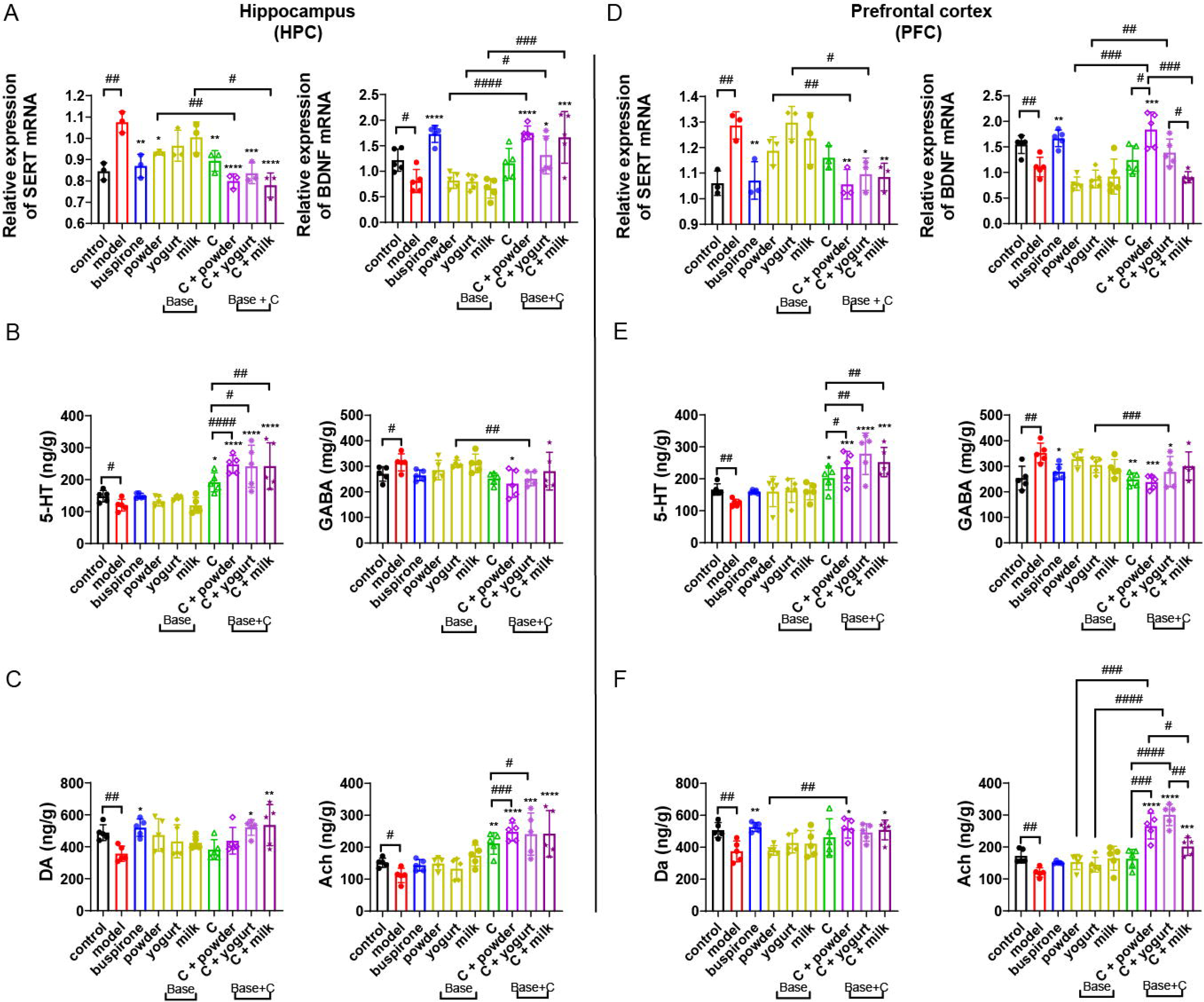
*C* re-balanced SERT and BDNF expressions in mouse-stress model. Panels **A**, **B**, and **C** present the findings pertaining to HPC assessment in mice, whereas panels **D**, **E**, and **F** depict the outcomes of PFC evaluation in mice. In the test results of the mouse HPC: (**A**) RT-qPCR analysis of SERT (left) and BDNF (right) in HPC tissue, results were normalized to β-actin, and presented as mean ± SEM, n = 3. (**B, C**) LC-MS/MS measued concentrations of 5-HT (**B**, left), GABA (**B**, right), DA (**C**, left), and ACh (**C**, right). In the test results of the mouse PFC: (**D**) RT-qPCR analysisof SERT (left) and BDNF (right) in HPC tissue. Data were normalized to β-actin and presented as mean ± SEM, n = 3. (**E, F**) LC- MS/MS measued concentrations of 5-HT (**E**, left), GABA (**E**, right), DA (**F**, left), and ACh (**F**, right). Data were presented as mean ± SEM, n = 5 per group. Significance was defined as *P* < 0.05 (\**P* < 0.05, \*\**P* < 0.01, \*\*\**P* < 0.001, \*\*\*\**P* < 0.01 vs. model group. #*P* < 0.05, ##*P* < 0.01, ###*P* < 0.01, ####*P* < 0.0001).

### SERT exhibits strong correlations with stress and cognitive-related indicators

In a randomized clinical trial, participants who underwent 9 months of meditation training exhibited a significant increase in serum BDNF levels, which correlated with reduced long-term cortisol levels and an increased volume of the dentate gyrus in the hippocampus^43^. Although direct clinical studies on SERT in the hippocampus are scarce, its function in the prefrontal cortex is strongly implicated in the pathogenesis of neuropsychiatric disorders, including depression^43^. Next, we conducted a comparative correlation analysis of various metrics between the human BO-stress model and mouse-stress model, with the latter further stratified into two brain regions, the HPC and the PFC (**Figure 6A**). Our findings revealed high correlations among BDNF, SERT, and neurotransmitters (5-HT, DA, and Ach) within the HPC and PFC of mice. However, no correlation was observed between HPC and PFC for GABA. The correlation between the metrics of the BO-stress and mouse-stress models was not significant. Notably, a significant correlation was observed between BO and HPC in terms of SERT expression, with a correlation coefficient of 0.640 (**Figure 6B**).

**Figure 6.**
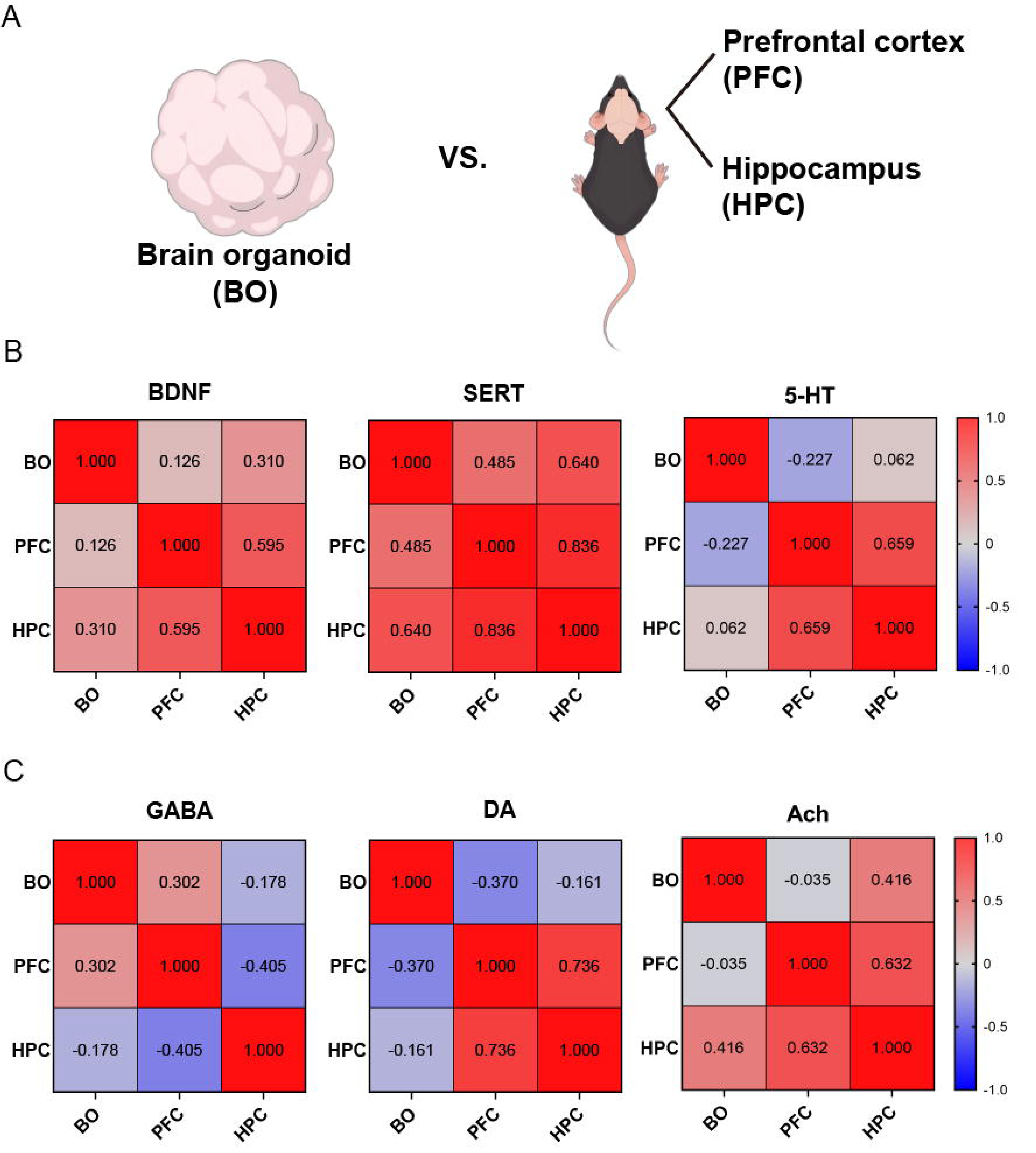
SERT exhibits strong correlations with stress and cognitive-related indicators. (**A**) Schematic illustration depicting the comparative correlation between BO and mouse-stress models. HPC and PFC of the mice were analyzed seperately. (**B, C**) Heatmaps illustrate correlation coefficients among indicators for individual mice, with color intensity indicating correlation strength:red for stronger, blue for weak correlations. Correlations in BDNF (**B**, left), SERT (**B**, middle), 5-HT (**B**, right), GABA (**C**, left), DA (**C**, middle), and ACh (**C**, right) between BO and mouse-stress models were analyzied.

Given the significant correlation observed between BO and SERT expression in the hippocampus, we proceeded to investigate the associations between hippocampal SERT levels and various other metrics in mice (**Supplementary** Figure 5A). Among all neurotransmitters, the concentration of SERT in HPC was negatively correlated with stress-like behaviors like EPM (r = -0.710, *P* = 0.002, **Supplementary** Figure 5C), as well as cognitive-related behaviors like recognition index in NOR (r = -0.684, *P* = 0.030, **Supplementary** Figure 5D) and PAT latency (r = -0.914, *P* < 0.001, **Supplementary** Figure 5E). The serum ACTH was positively correlated with SERT (r = 0.935, *P* < 0.001, **Supplementary** Figure 5F). The serum corticosterone was positively correlated with SERT (r = 0.881, *P* < 0.001, **Supplementary** Figure 5G). The same correlation was found between 5-HT and SERT (r = 0.816, *P* = 0.004, **Supplementary** Figure 5H). The BDNF was positively correlated with SERT (r = -0.843, *P* = 0.021, **Supplementary** Figure 5L). This suggests that SERT is highly correlated with other markers in the mouse hippocampus and may serve as a key target for regulating the balance between stress and cognitive.

## Discussion

In this study, we first examed the expression of neurontransmitters and SERT/BDNF, after the addition of *Th, WP* and *Th+WP* in BO-stress model. We found that, *Th*+*WP*suppress the expression of SERT, while enhanced the expression of BDNF. We then selected the (*Th+WP*)-low dose (at 85 + 200 mg/mL, termed as the ***C***), or ***C*** supplemented with powder, milk, and yogurt (the vehicles), for efficacy evaluation in the mouse-stress model, and the results demonstrated stress-reducing and cognitive-enhancing effects. We compared the differences between BO and various indicators in mice, revealing interspecies variations in these metrics. Notably, SERT expression exhibited a positive correlation between humans and mice, with a correlation coefficient of 0.640 (**Figure 6B**). We then conducted a correlation analysis between SERT expression in the mouse hippocampus (HPC) and other related indicators. The results indicated that mouse stress and cognitive behavior, as well as hippocampal neurotransmitters, were closely linked. This suggests that SERT may be a key protein explaining why the combination of ***C*** with different bases in the mouse- stress model can both alleviate stress and enhance cognitive.

The BO represent a groundbreaking *in vitro* modeling approach that emulates structures and functions of the human brain, effectively bypassing the species-specific disparities inherent in rodent models. Up to now, scientists have achieved notable progress in establishing diverse models utilizing BOs^44^: Liu Y. et al (in 2008) employed BOs to clarify the mechanistic role of GABA-ergic inhibitory neurons in depression with suicidal ideation^45^. In addition, Xiang Y. et al in 2001 has developed BOs specific to the spinal trigeminal nucleus (SpV) and successfully constructed an *in vitro* model of nucleus-related brain interconnections^46^. However, research on the enhancement of BDNF through natural extracts and the reduction of SERT expression *in vitro* remains remains largely unexplored.

In mice study, the EOP creates standardized environments for mice, inducing stress-like behavior by replicating natural challenges^47^. Foot shock stress, while commonly used to induce stress in mice, has inconsistent procedures and lead to anhedonia and learned helplessness^48,49^. In addition, we employed OFT, EPM and LDB tests to evaluate stress levels in mice. Interestingly, the EPM and LDB tests demonstrated greater sensitivity in detecting anxiolytic effects compared to the OFT, aligning with findings from previous research^50^. Our previous study also showed that OFT results could be influenced by the administration method, particularly oral administration, potentially leading to depression and stress-like behaviors that could interfere with experimental outcomes^51^. In this study, we administered powder, yogurt, milk, and the ***C*** to the mice via intragastric gavage, which inevitably affected OFT results (**Figure 3C** and **Supplementary** Figure 5B). To ensure a comprehensive assessment, additional stress-related experiments are necessary.

Disruptions in stress regulation are frequently linked to dysfunctional neurotransmitter systems^52^. In primates, the PFC crucially regulates stress through advanced strategies for modifying and coping with stress experiences^53^. Hippocampal dysfunction is intimately associated with the pathogenesis and progression of a range of neurodegenerative disorders, such as Alzheimer’s Disease (AD), epilepsy, depression, and anxiety^54^. Based on these, we measured the concentrations of four neurotransmitters in PFC and HPC of mice. Research involving chronic social defeat stress and chronic restraint stress models in mice has documented a reduction in 5-HT level within PFC, corroborating our findings in this study^55^. Compared with the non-chronic mild stress (non-CMS) group, the CMS mice had significantly lower GABA level in PFC^56^. Upregulated GABAAα2 and GABAAγ2 expression existed in the brain PFC of restraint stress mice^57^. Using a microdialysis study, Saitoh A. et al in 2018 found increased extracellular GABA level of HPC in mice with veratrine-induced stress-like behaviors^58^. The role of DA in stress and cognitive, particularly within the PFC and HPC, remains inadequately understood. DA is critical for cognitive processes reliant on prefrontal function, as exemplified by observations in Parkinson’s disease^59^. The concentration of DA in PFC was usually reduced in stress-like behavior mice^60^. ACh functions as a key neurotransmitter within theHPC, playing a role in modulating cognitive processes. In individuals afflicted with AD, reduction in ACh levels within the hippocampal region is observed, which correlates strongly with cognitive deficits^61^.

Previous studies have shown reduced BDNF protein expression in the HPC induced by chronic stress^62^, which aligns with our results. The HPC, known for its high sensitivity to internal stressors, plays a vital role in regulating emotional responses. Moreover, it’s the primary site where BDNF exerts its activity^63^. SERT principally mediates the reuptake of 5-HT, thereby modulating its concentration within the synaptic cleft and influencing neurotransmission. In HPC, alterations in SERT activity are linked to mood regulation, cognitive formation, and cognitive deficits^38^. We innovatively analyzed the correlation of various metrics between BO and mouse-stress models. We discovered a correlation in SERT expression between human and mouse brain, suggesting its potential as a molecular target for evaluating preclinical indicators and elucidating its underlying mechanisms.

Investigations have underscored the pivotal role of low-grade inflammation in the onset of stress, frequently associated with increased levels of pro-inflammatory cytokines in the brain^64^. Furthermore, inflammation can lead to a shift in tryptophan metabolism from the 5-HT pathway to the kynurenine pathway^65^. Nevertheless, we found no significant differences between the model and control, potentially due to the specific brain region selected for IF staining and the timing of sample collection. Furthermore, activated microglia can manifest distinct phenotypes, including M1, which contribute to inflammation, and M2, which have anti-inflammatory effects^66^. It’s possible that more precise and meaningful results could be obtained if we separately assessed the M1 and M2 microglia phenotypes.

## Conclusion

Utilizing the BO-stress model, we demonstrated that *WP+Th* reduced the expression of the SERT while upregulated BDNF. Further researches revealed *WP* (85 mg/mL) + *Th* (200 mg/mL), designated as the ***C***, significantly improved the behavioral performance of mice in stress and cognitive assays. In addition, ***C*** was demonstrated to rebalance neurotransmitter, as well as SERT and BDNF. The synergistic effects of *Th* and *WP* suggest a potential for stress mitigation and the elevation of neurotrophic factor levels. We assessed the correlation of various metrics between BO and mouse stress models, identifying a shared correlation of SERT. Notably, SERT expression in the mouse HPC was associated with stress, cognitive performance, and neurotransmitter levels.

## Funding information

This study was supported by: 1. National Key Research and Development Program of China (2021YFF0702000); 2. National Natural Science Foundation of China (82071349); 3. Sichuan Science and Technology Program (2025ZNSFSC0703); 4. West China Hospital of Sichuan University Discipline Excellence Development 1 3 5 Engineering Project (ZYYC08005 and ZYJC18041); 5. National Center of Technology Innovation for Dairy (2023-JSGG-13).

## Conflict of interest

Xiuzhen Jia, Zifu Zhao, Jingyu Hao, and Jian He are affiliated with Inner Mongolia Yili Industrial Group Co., Ltd. All other authors declare no competing financial interests or commercial relationships that could be construed as a potential conflict of interest.

## Authors contribution statements

Qixing Zhong, Qinxi Li, and Xiuzhen Jia conducted all the experimental procedures and drafted the manuscript. Linxue Hu, Yingqian Zhang, Jingyang Zu, Yao He and Xiaojie Li helped to conducted the experimental procedures. Yu Wang, Haotian Feng, and Jingyu Hao prepared the nutrient combinations, while Zifu Zhao and Jian He calculated their concentrations. Zhihui Zhong contributed to the conception and design of the study. The final manuscript was carefully reviewed and approved by all authors.

## Acknowledgement

The authors would like to thank Sichuan Junhui Biotechnology Co., Ltd. and Singapore HumanSim Pte. Ltd. for providing the experimental platforms for this study.

## Data availability statement

The authors confirm that the data supporting the findings of this study are available within the article and its supplementary materials.

## Notes

### Competing Interest Statement

The authors have declared no competing interest.

