## Supplementary materials for "Synergistic neuroprotective and cognitive-enhancing effects of Walnut Peptide and Theanine in human brain organoid and mouse stress models"

**Supplementary Tables**

**Supplementary Table 1.** The chromatographic gradient of mobile phase (A: water, and B, acetonitrile)

|  | **Flow rate (mL/min)** | **A (%)** | **B (%)** |
| --- | --- | --- | --- |
| **Initial** | 0.6 | 90 | 10.0 |
| **0.30** | 0.6 | 90 | 10.0 |
| **2.00** | 0.6 | 10.0 | 90.0 |
| **2.50** | 0.6 | 10.0 | 90.0 |
| **2.60** | 0.6 | 90.0 | 10.0 |
| **3.50** | 0.6 | 90.0 | 10.0 |

**Supplementary Table 2.** Antibodies used for IF staining.

|  | **Antibody name** | **Host Species** | **Dilution** | **Catalogue**  **number** | **Manufacturer** |
| --- | --- | --- | --- | --- | --- |
| **Primary**  **antibody** | PAX6 | Rabbit | 1:100 | ab195045 | abcam |
|  | EN1 | Rabbit | 1:100 | ab195045 | Invitrogen |
|  | GFAP | Rabbit | 1:100 | ab68428 | abcam |
|  | MAP2 | Rabbit | 1:100 | 13-1500 | CST |
|  | TUJ1 | Rabbit | 1:100 | 4466S | CST |
|  | BDNF | Rabbit | 1:100 | ab108319 | abcam |
|  | Serotonin transporter | Rabbit | 1:100 | PA5-80032 | thermofisher |
| **Secondary**  **antibody** | FITC labeled (Green) | Goat anti rabbit | 1:200 | GB22303 | Servicebio |
|  | CY3 labeled (Red) | Goat anti mouse | 1:200 | GB21301 | Servicebio |

**Supplementary Table 3.** Primers used for RT-qPCR analysis

| **Primers** | **Forward sequence (5’→3’)** | **Reverse sequence (5’→3’)** |
| --- | --- | --- |
| **Host Species: mouse** | | |
| β-actin | CCACCATGTACCCAGGCATT | CAGCTCAGTAACAGTCCGCC |
| BDNF | TCCGGGTTGGTATACTGGGTT | GCCTTGTCCGTGGACGTTT |
| SERT | GCATCCACATTCTTTGCCATCAT | TGATGACCACGATGAGCACAAAC |
| **Host Species: human** | | |
| GAPDH | TGACATCAAGAAGGTGGTGAAGCAGG | GCGTCAAAGGTGGAGGAGTGGGT |
| OCT4 | CTGGGTTGATCCTCGGACCT | CCATCGGAGTTGCTCTCCA |
| Nanog | GATTTGTGGGCCTGAAGAAA | CAGATCCATGGAGGAAGGAA |
| TUBB3 | AAGACGACGAGGAGGAGTCG | GGGGTTTAGACACTGCTGGC |
| EN1 | CGCCCAGTTTCGTTTTCGTT | GCAGAACAGACAGACCGACA |
| GFAP | GCACGCAGTATGAGGCAATG | TAGTCGTTGGCTTCGTGCTT |
| PAX6 | CTGAAGCGGAAGCTGCAAAG | TTGCTGGCCTGTCTTCTCTG |
| RBFOX3 | ACCCTACAGAGAAGCAGCAG | CGAATTGCCCGAACATTTGC |
| MAP2 | GGAAAACCACAGCAGCAGGT | CGAGGAGGGAGAATGGAGGA |
| SOX2 | TTGCGTGAGTGTGGATGGGATTGGTG | GGGAAATGGGAGGGGTGCAAAAGAGG |
| BDNF | CAGGGGCATAGACAAAAGGC | TCCTTATGAATCGCCAGCCA |
| SERT | GCAGGGACGTGAAGGAA | GTACCACCGAAATGGATG |

**Supplementary Figures**

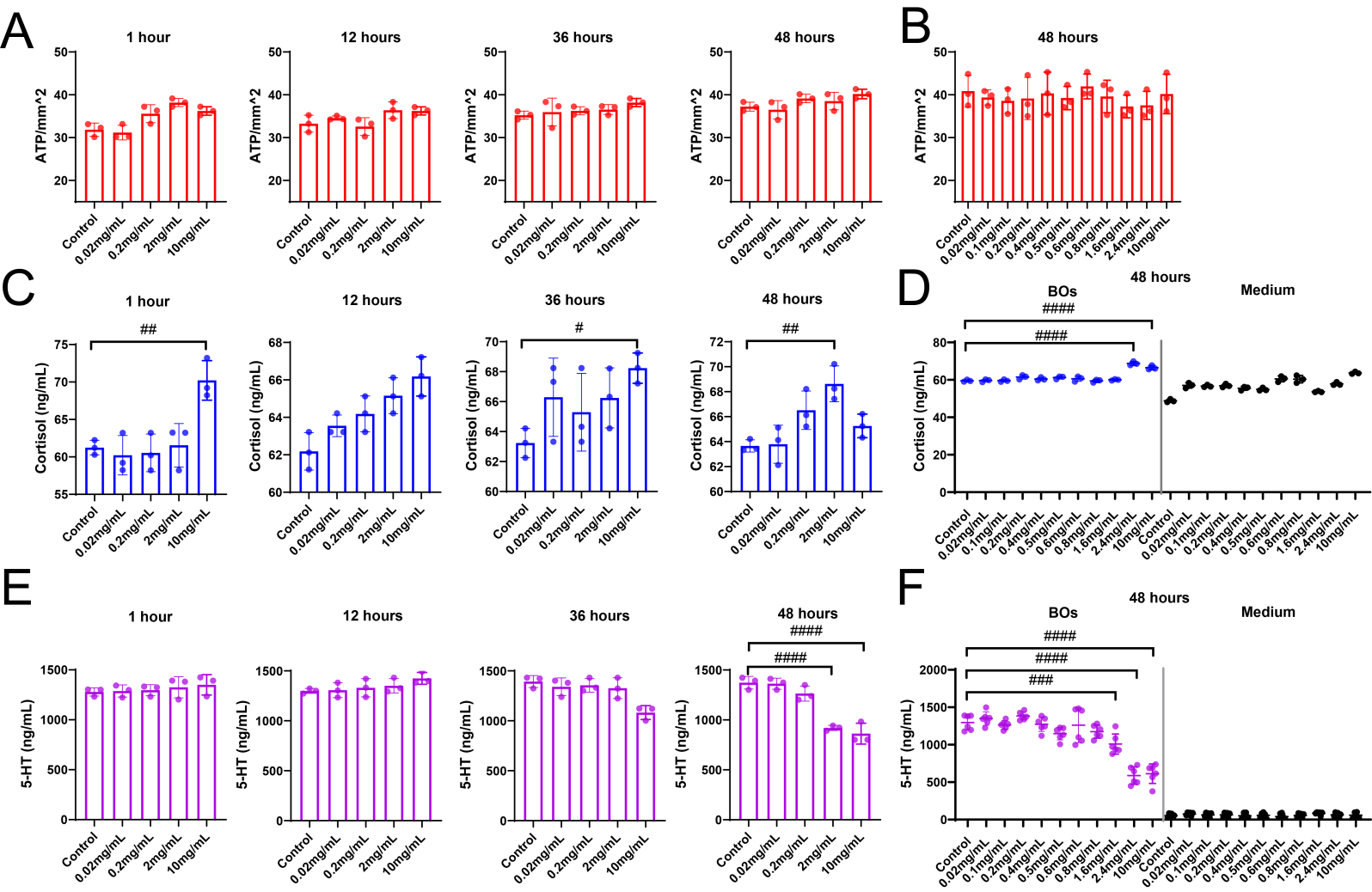

**Supplementary Figure 1.**

The concentrations of ATP (**A**, **B**), cortisol (**C**, **D**) and 5-HT (**E**, **F**) in the BOs after mCPP addition on day 80 and in their culture medium, were measured by ELISA. The data were presented as mean ± SEM. **A**-**D**: n = 3, **E**-**F**: n = 6. Significance was defined as *P* < 0.05 (#*P* < 0.05, ##*P* < 0.01, ###*P* < 0.001, ####*P* < 0.0001).

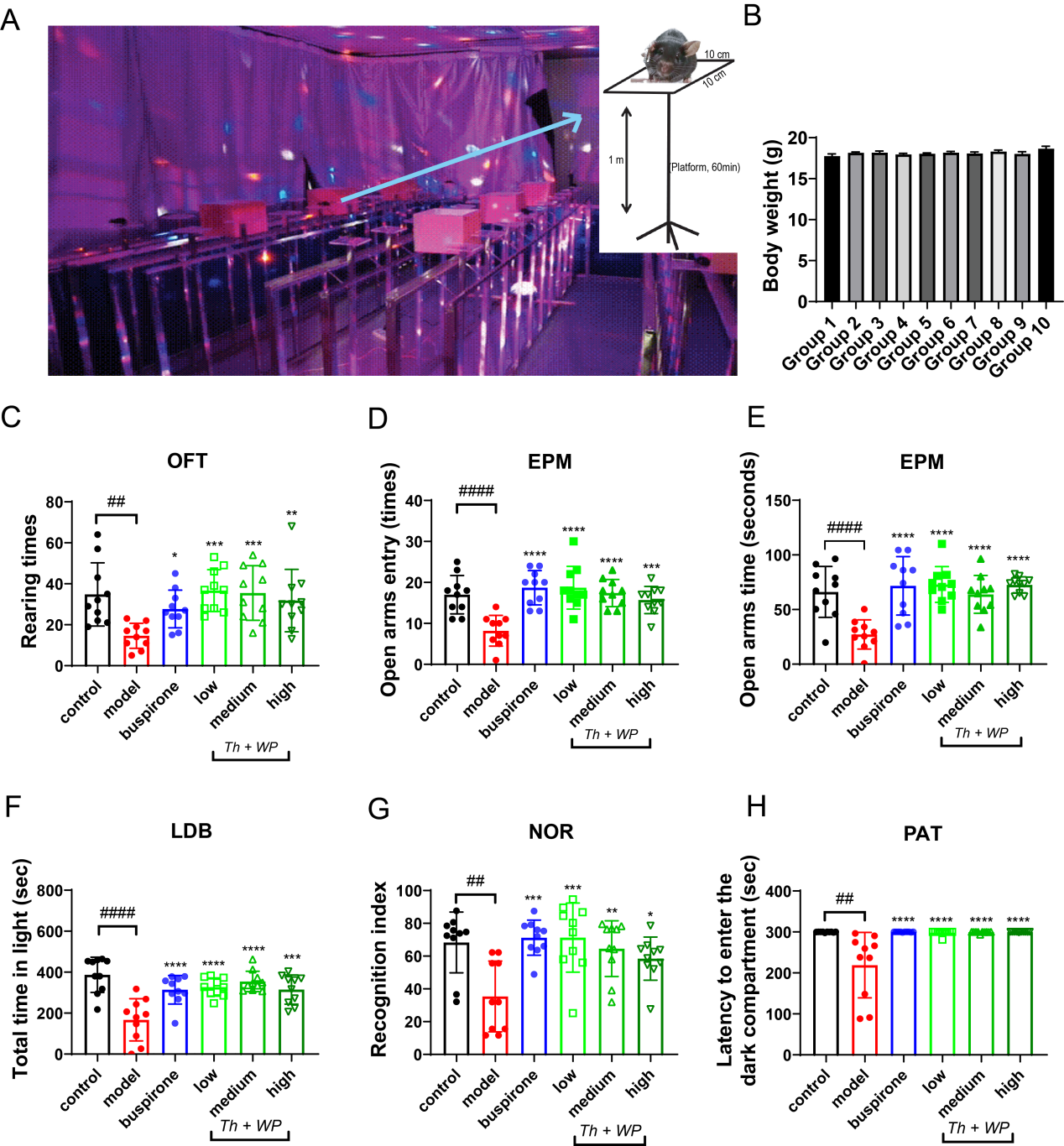

**Supplementary Figure 2.**

(**A**) EOP and mouse-stress model: We set individual high platforms sequentially within the stress-inducing environment, housing 5 mice in isolation per row, culminating in a total of three rows. (**B**) On day 1, mice were randomly assigned to groups based on body weight, with weights recorded. Data are presented as mean ± SEM. OFT performed on day 30. Rearing times (**C**). EPM performed on day 32. Changes of open arms entry (**D**) and open arms time (**E**). LDB performed on day 35. Total time in light (**F**). Schematic of novel object recognition test. Recognition index analyzed by NOR in each group (**G**). On day 36, the avoidance test was conducted. Schematic of the PAT (**H**). The data were presented as mean ± SEM, with n = 10 in each group. Significance was defined as *P* < 0.05 (**P* < 0.05, ***P* < 0.01, ****P* < 0.001, *****P* < 0.01 vs. model group. ##*P* < 0.01, ####*P* < 0.0001).

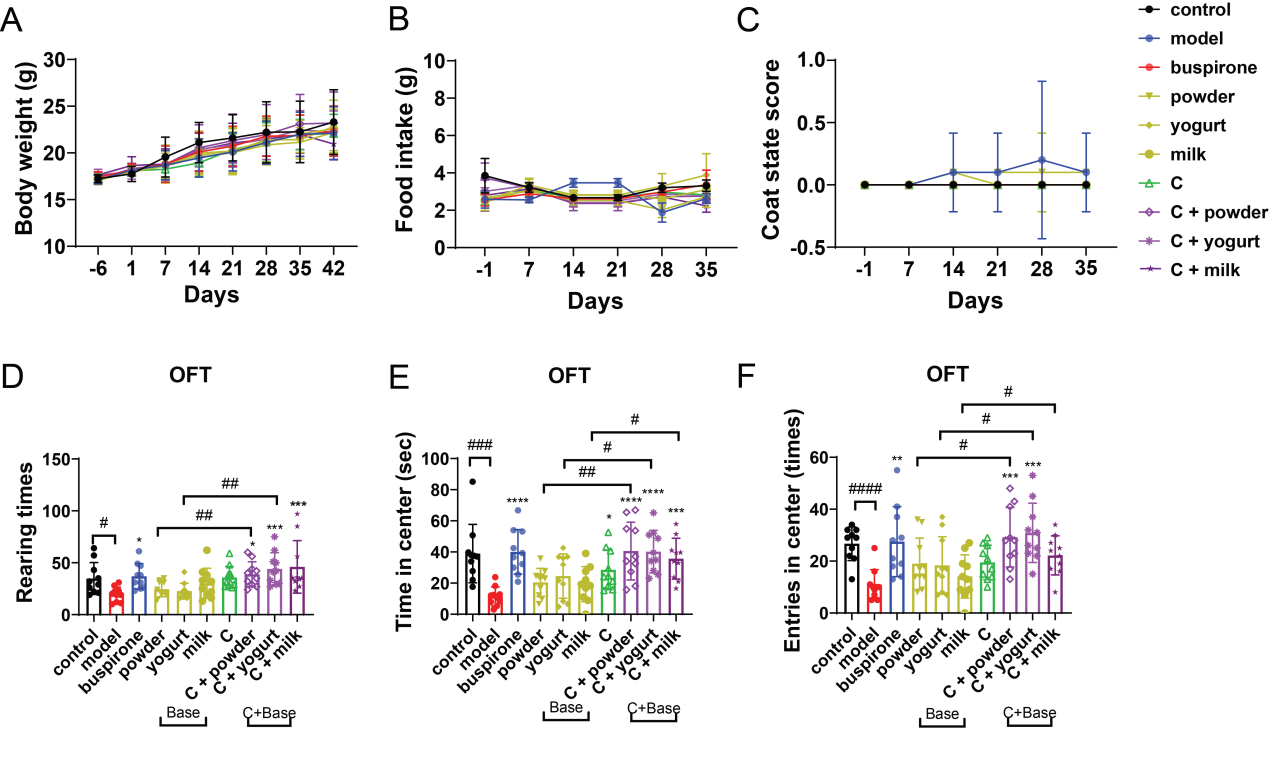

**Supplementary Figure 3.**

Weekly assessments of body weight (**A**), food consumption (**B**), and coat condition scores (**C**) were conducted. Data were presented as mean ± SEM, n=10. Repeated measures ANOVA was used to identify inter-group variations. Post-hoc Bonferroni tests were applied to significant results. No significant differences were detected across the three parameters in all groups. On day 14, OFT analyses included rearing frequency (**D**), time spent in the inner zone (**E**), and central entries (**F**) for each group. Data were presented as mean ± SEM, n=10. Unpaired t-tests for pairwise comparisons and one-way ANOVA with Bonferroni post hoc tests were used for multiple group comparisons. Significance was defined as *P* < 0.05 (**P* < 0.05, ***P* < 0.01, ****P* < 0.001, *****P* < 0.01 vs. model group. #*P* < 0.05, ##*P* < 0.01, ###*P* < 0.001, ####*P* < 0.0001).
.

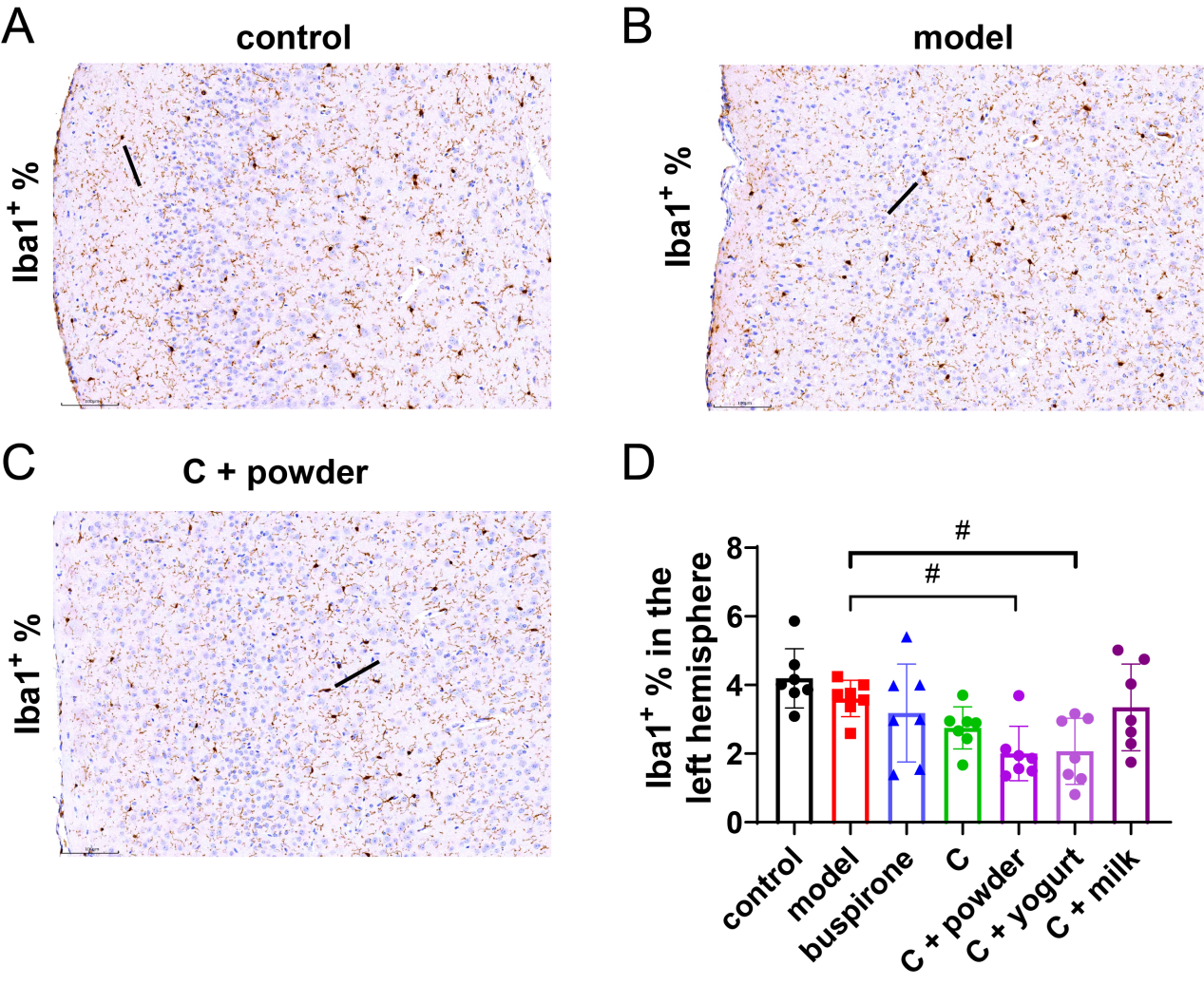

**Supplementary Figure 4.**

In each group, the left hemisphere was extracted and immunohistochemical staining was employed to evaluate the expression of Iba1. Three representative images, denoted as **A, B,** and **C**, correspond to the three IF groups. The black short lines marked the bushy Iba1^+^ Cells. **D.** The variations in the percentage of astrocytes showing positive expression between different groups. Data were presented as mean ± SEM, n = 7. Significance was determined at *P* < 0.05 (#*P* < 0.05).

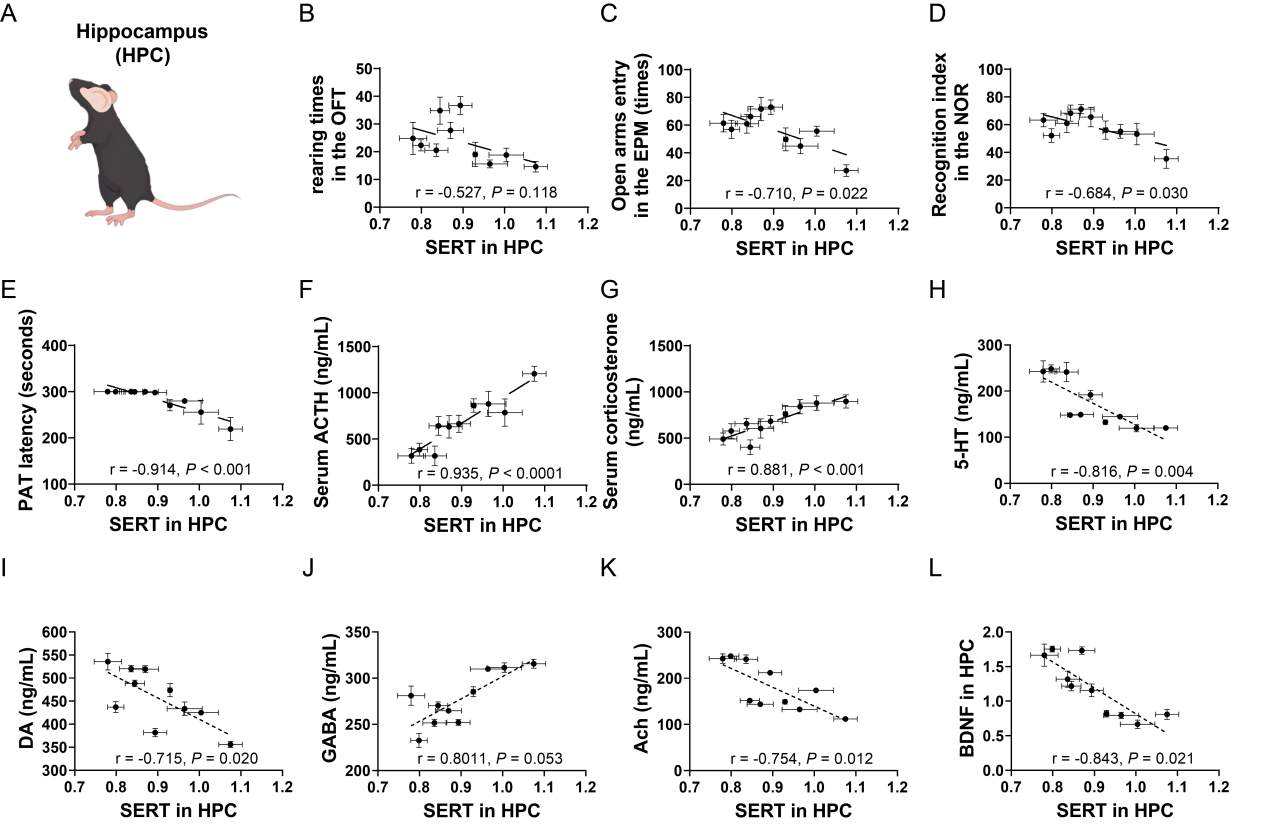

**Supplementary Figure 5.**

(**A**)**.** Illustrative schematic of the mouse hippocampal area. In OFT, 'Rearing times' served as the index; in EPM, 'Open arms entry' was selected; 'Total time in light' was the metric for both LDB and NOR experiments. Finally, in the PAT experiment, the 'Latency to enter the light compartment' was the chosen metric. We assessed correlations between SERT mRNA levels in the HPC and rearing frequency in the OFT (**B**), open arm entries in the EPM test (**C**), recognition indices in the NOR test (**D**). PAT latencies (**E**), serum ACTH levels (**F**), corticosterone levels (**G**), DA(**I**), GABA(**J**), Ach(**K**), and BDNF(**L**).
